# Competitive catabolism in systemic metabolic homeostasis

**DOI:** 10.1101/2025.10.29.684184

**Authors:** Daniel R. Weilandt, Won Dong Lee, Michael R. MacArthur, Lingfan Liang, Corey Holman, Qingwei Chu, Alexis J. Cowan, Joseph A. Baur, Ned S. Wingreen, Joshua D. Rabinowitz

## Abstract

Systemic metabolic homeostasis maintains circulating nutrient concentrations within physiological ranges. Insulin is central to this process, lowering circulating levels of glucose, free fatty acids, and ketones. Yet, how simultaneous homeostasis of these nutrients is achieved remains unclear. Here we develop a differential equation model of fasting metabolic homeostasis. Grounded in mass action kinetics, this multi-nutrient model reveals how a fixed energy demand naturally leads to competition between major circulating nutrients for oxidation (‘competitive catabolism’). Perturbative nutrient infusions confirm this emergent behavior. The multi-nutrient model predicts that insulin promotes fasting glucose homeostasis primarily indirectly by slowing lipolysis. It further identifies a physiological circuit by which obesity causes diabetes: Increased fat mass promotes lipolysis, releasing fatty acids into circulation that compete with glucose for oxidation, elevating glucose. Resulting hyperinsulinemia restores proper lipid catabolic flux but not euglycemia. Thus, quantitative modeling reveals a physiological homeostatic circuit through which obesity causes type 2 diabetes.

## Introduction

In mammals, organismal metabolic homeostasis requires insulin, which is secreted by pancreatic β-cells in response to elevated blood glucose levels.^1^ Insulin increases glucose disposal, establishing a feedback loop that maintains glucose homeostasis. Loss of this feedback loop due to autoimmune β-cell destruction leads to type 1 diabetes.^2^ Deficient response to insulin (‘insulin resistance’) results in type 2 diabetes^3^, with the primary risk factor for type 2 diabetes being obesity^4^.

Insulin acts on multiple tissues to lower blood glucose^5^, as well as ketones^6^ and free fatty acids^7^. In muscle, insulin promotes GLUT4 translocation and hence glucose uptake.^8^ In the liver, it triggers phosphorylation of glycogen synthase and glycogen phosphorylase, favoring glucose storage as glycogen.^9^ It also promotes transcription of lipogenic genes, driving fat synthesis.^10^ In adipose, insulin lowers cyclic AMP levels to suppress lipolysis, promoting fat storage.^11^

Beyond these direct actions, insulin also impacts metabolic fluxes indirectly through interactions spanning different nutrients and tissues. For example, insulin suppresses gluconeogenesis and ketogenesis in part by lowering circulating glycerol and free fatty acids.^12^ In this circuit, insulin’s direct action on adipose to suppress lipolysis indirectly slows hepatic pathways making glucose and ketones. Similarly, insulin indirectly promotes lactate burning in muscle by acting directly upon adipose to lower circulating free fatty acids.^13^

Relative to linear metabolic or signaling pathways, these multi-nutrient homeostatic circuits are more difficult to dissect using experiments alone. Mechanistic quantitative models provide a complementary approach to disentangle how metabolites, hormones, and tissues coordinate metabolic homeostasis. Existing quantitative models tend to simulate insulin acting on individual metabolites to mimic postprandial dynamics, without accounting for cell-mediated metabolite-metabolite interactions.^14,15^ These models successfully mirror temporal changes of metabolites following a meal^16–18^ or glucose challenge^19–22^, but fail to explain how the basal homeostatic state is established and maintained. Through manipulation of the models’ parameter values, they can also mimic obesity or diabetes. However, this requires external alteration of the model to change factors like insulin sensitivity or β-cell glucose set points, as these models do not capture an intrinsic relationship between obesity and diabetes.

Here, we develop a multi-nutrient model of fasted metabolic homeostasis that includes both insulin-metabolite and metabolite-metabolite interactions. Both the model and perturbative infusion experiments support a key role in systemic metabolic homeostasis for competition among major nutrients to enter TCA cycle metabolism and be oxidized (competitive catabolism). The model further reveals an intrinsic physiological connection between obesity and hyperglycemia: By mass action, elevated fat mass promotes lipolysis. Due to competitive catabolism, lipolysis tends to elevate glucose, increasing insulin. Elevated insulin feedback inhibits lipolysis at the expense of persistent hyperglycemia and hyperinsulinemia.

## Results

### Multi-nutrient model of organismal metabolism

We set out to develop a quantitative model of fasting mammalian organismal metabolism, incorporating insulin and four major circulating carbon-carrier fuels (glucose, lactate, ketone bodies, and fatty acids; Figure 1A). For each, we describe their rate of change in the form of a differential equation:

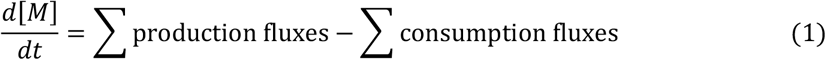

**Figure 1:**
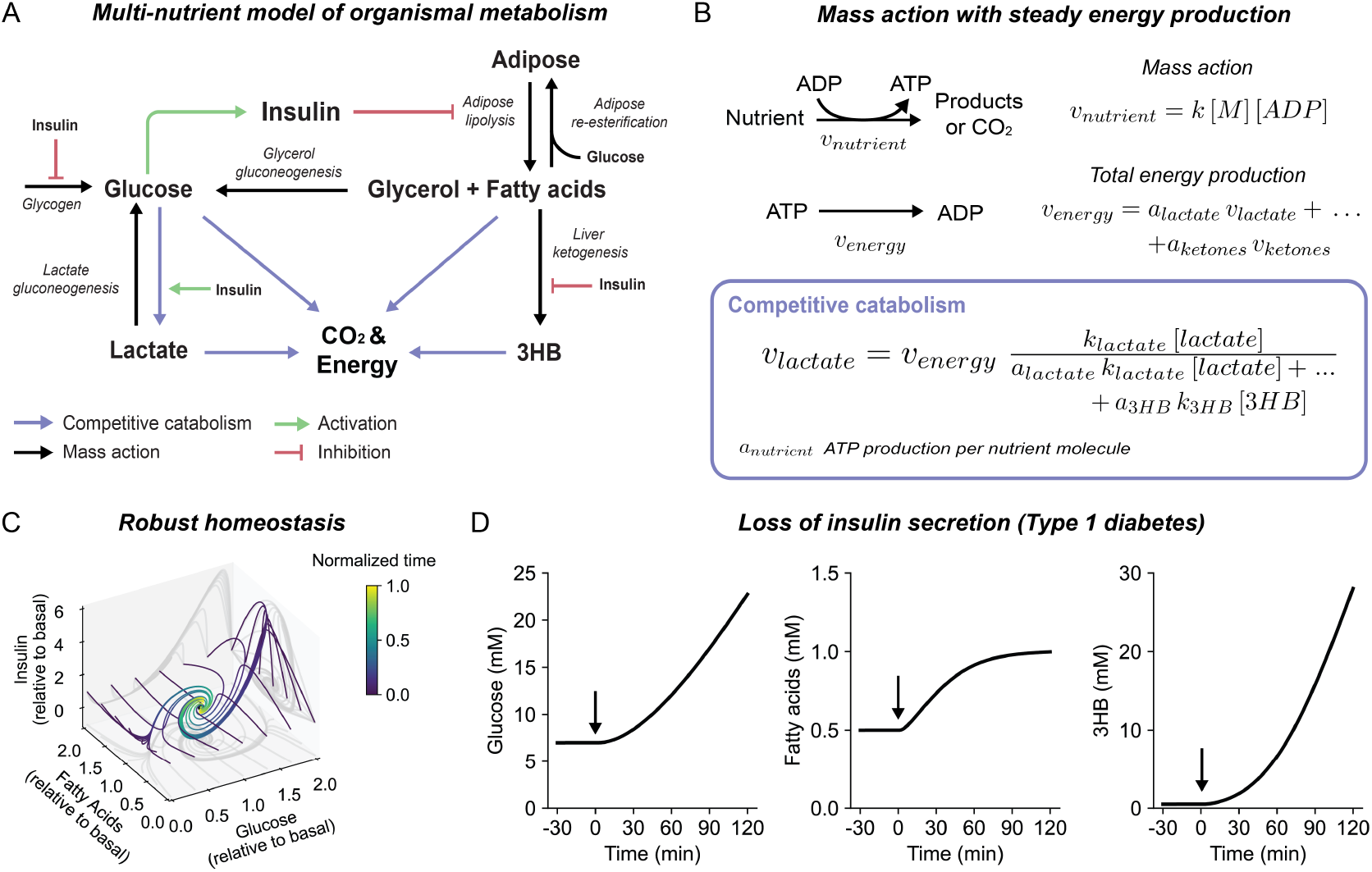
A multi-nutrient model of fasted organismal metabolism. A) Model schematic. B) Mass action kinetics under the constraint of stable energy expenditure leads to the competitive catabolism (see text Eq. 3). The competitive catabolism equation describes, as a function of circulating nutrient levels, the rates of glucose, lactate, fatty acid, and ketone oxidation (blue arrows in the model in panel A). C) The multi-nutrient model converges to a stable steady state. Lines show trajectories of glucose, fatty acids, and insulin as the network returns to steady state from different perturbations. D) Loss of insulin secretion yields an uncontrolled rise in glucose and ketones, mimicking type 1 diabetic ketoacidosis. 3HB denotes the ketone body 3-hydroxybutyrate.

Insulin production and consumption are modeled as in prior literature^18,72^: Saturable, ultrasensitive glucose-triggered production and mass action consumption. Metabolic reactions are glucose production by glycogen breakdown and by gluconeogenesis (which consumes lactate), glucose consumption by glycolysis (which produces lactate), free fatty acid production by lipolysis, and free fatty acid consumption by reesterification and by ketogenesis (which produces ketone bodies). These metabolic reactions are modeled by simple mass action kinetics (Table S1). Insulin, with a delay from secretion to peak action, suppresses ketogenesis and lipolysis and induces glycolysis.^18^

In the fasted state, the primary fate of circulating nutrients is oxidation to generate ATP. In prior work, we have found that such oxidation generally follows mass action kinetics:^23^

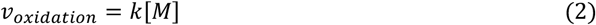

While simple mass action is a good approximation for individual minor nutrients like specific amino acids, generalization to all nutrients would predict that whole-body ATP production (calorie burning) is a linear function of circulating substrate availability. In contrast, ATP production rate is thought to be mainly controlled by demand, triggered by activities like moving or thinking, not circulating metabolite levels.^24,25^ Assuming that demand is insensitive to circulating metabolite levels, and steady-state ATP production must equal demand, we solved for energy production fluxes (glycolysis and oxidation of each nutrient) (Figure 1B):

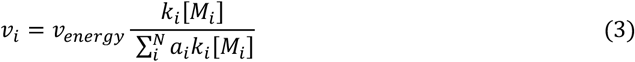

Here, *v*_*energy*_ is whole body energy expenditure, *k*_*i*_ is the rate constant reflecting the propensity to catabolize nutrient *i*, and *a*_*i*_ is the ATP yield from such catabolism. The resulting equation reveals competitive catabolism: The catabolism of each nutrient is suppressed by every other nutrient, to preserve constant ATP flux (Figure S1A). Structurally, the competitive catabolism equation resembles competitive Michaelis-Menten kinetics, except (i) collectively nutrients must always fill energy demand (*v*_*energy*_) and (ii) the competitive impact of each nutrient depends not only on its mass action kinetics (*k*_*i*_[*M*_*i*_]) but also the number of ATP (*a*_*i*_) generated in its catabolism.

### Model parameters

Based on mass action and competitive catabolism, we assembled a multi-nutrient model of fasting metabolic dynamics. The model comprises 24 quantitative parameters, largely mass-action rate constants, as well as affinity constants characterizing insulin action on different pathways (*K*_*a*_, *K*_*i*_). Because the model is based on mass action kinetics, the number of parameters is substantially less than prior metabolic modeling efforts of similar complexity. Combined with greater availability of experimental data including metabolite concentrations and fluxes, this allowed rigorous determination of parameter values to obtain a predictive model.

Specifically, four parameters set the turnover time scales of glucose, lactate, fatty acids, and ketones, respectively, and were experimentally determined by steadily infusing the ^13^C-labeled form of each fuel and measuring the time it takes to reach half of the steady-state labeling in the circulation^13^. Five parameters characterizing insulin dynamics were taken from literature while five determining its action were chosen to reflect physiological observations. As insulin increases, it first suppresses ketogenesis, then suppresses lipolysis, and finally directly impacts glucose metabolism (Note: smaller affinity constant *K* implies greater sensitivity to small quantities of insulin; Table S2):^26–28^

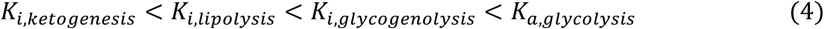

This left ten parameters to describe a total of ten fluxes; therefore, these parameters could be directly calculated algebraically by introducing experimentally measured steady-state fluxes and concentrations as constraints (Figure S1B, Tables S2,3).

The resulting parameterized model desirably converged to a stable steady state and was robust to perturbations in any of the four nutrients and insulin (Figure 1C and S1C). Moreover, the model was insulin-dependent in a manner that simulates the pathophysiology of type 1 diabetes: Upon removal of insulin, blood glucose, fatty acids, and ketones increase in an uncontrolled manner, mimicking diabetic ketoacidosis (Figure 1D). Thus, three key ingredients – mass action kinetics, competitive catabolism, and insulin action – are sufficient to capture key aspects of mammalian metabolic homeostasis.

### Competitive catabolism

Competitive catabolism involves two core tenets: (i) Each nutrient drives its own oxidation via mass action (saturating as energy production from this substrate approaches total energy expenditure), and (ii) energy expenditure is unresponsive to nutrient levels. To test these tenets, we infused catheterized fasted mice with ^13^C-labeled glucose, lactate, and 3-hydroxybutyrate (3HB), at either a low rate (minimally perturbative, to probe baseline metabolism) or a high rate (intentionally perturbative, aiming to increase the concentration of the infused metabolite to induce its catabolism) (Figure 2A). During the infusions, the mice were housed in metabolic cages and total exhaled CO_2_, ^13^CO_2_ enrichment, and energy expenditure (based on oxygen consumption) were measured. Together with measurements of circulating substrate ^13^C-labeling, these data enable quantitation of fuel-specific catabolic fluxes.

**Figure 2:**
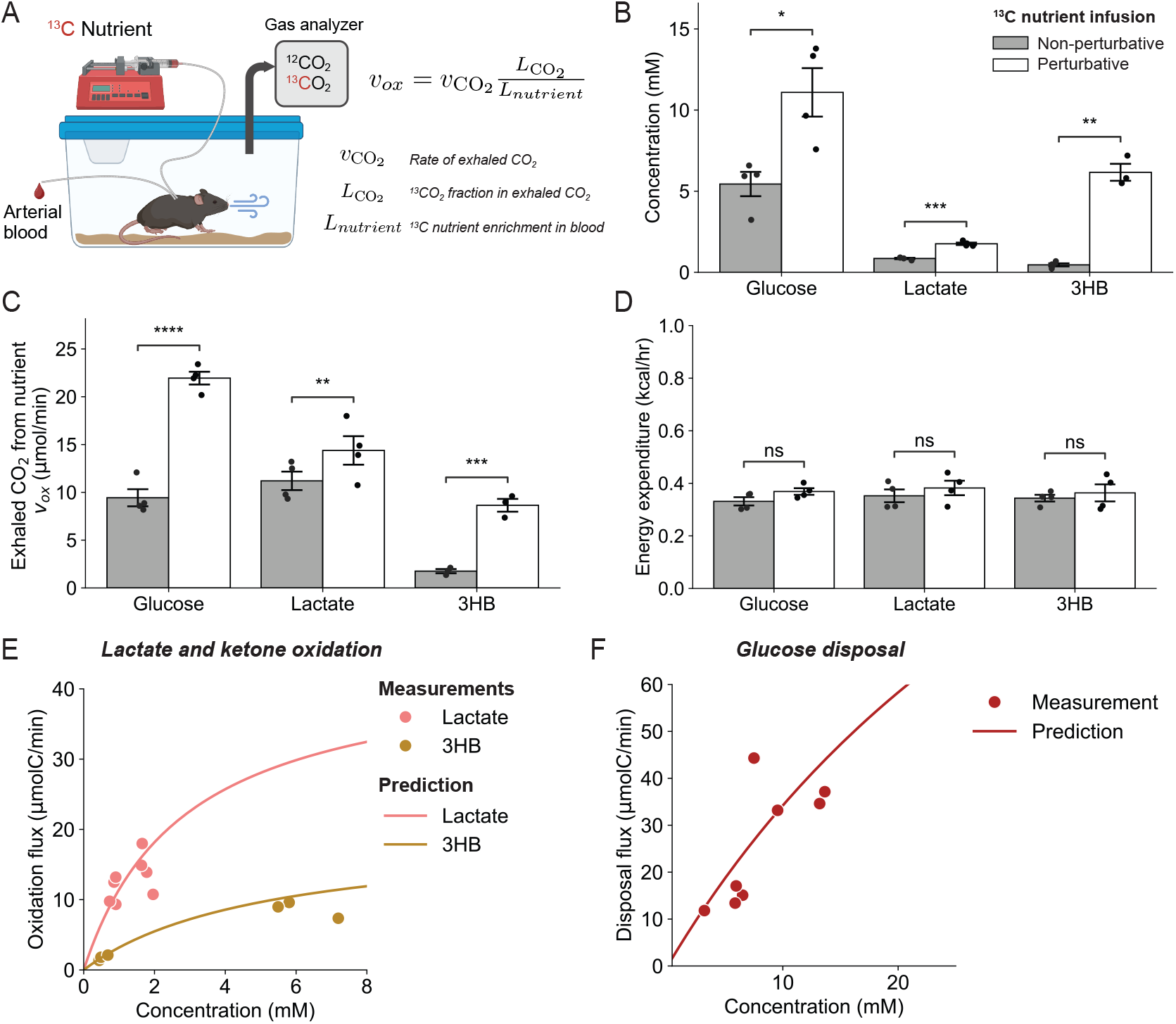
Experimental evidence for competitive catabolism. A) Experimental setup: Doubly catheterized mice are infused within a metabolic cage with ^13^C-labeled nutrients at either an intentionally perturbative or minimally perturbative rate. Concentrations and ^13^C-labeling of both arterial blood metabolites and exhaled gases (CO_2_, O_2_) are measured, enabling determination of whole-body energy expenditure and catabolic fluxes. B) Perturbative infusions effectively raise the arterial concentrations of the infused nutrient (n=3-4 mice per nutrient infusion). C) Perturbative infusions increase the infused nutrient’s oxidation flux. Measured exhaled ^13^CO_2_ from the infused ^13^C-metabolite (glucose, lactate, and 3HB) is reported normalized to the metabolite’s circulating labeling fraction (n=3-4 mice per nutrient infusion). D) Perturbative infusions do not alter whole body energy expenditure as measured by indirect calorimetry (n=3-4 mice per nutrient infusion). E) Measured oxidation flux (data points) matches the competitive catabolism equation (lines), for lactate and 3HB (n=3-4 mice per nutrient infusion). F) Same for disposal flux of glucose (glucose oxidation + glycolysis) (n=3-4 mice per nutrient infusion). 3HB, 3-hydroxybutyrate. ns, not significant. *p < 0.05, **p < 0.01, ***p < 0.001, and ****p < 0.0001 by two-sample t test. Data are mean ± SEM.

Perturbative infusions consistently elevated the infused metabolite’s circulating concentration to a new steady state (Figure 2B). For glucose and 3HB, their oxidation rates increased between 2 and 5-fold, roughly proportional to the increase in circulating concentration (mass action). In contrast, lactate infusion led to only a modest increase in total lactate oxidation rate. As lactate is a major fasted-state fuel, this modest increase in oxidation rate can be explained by lactate-driven energy production approaching total energy expenditure (saturation limiting mass action). In all cases, whole body energy production was invariant (Figure 2C). Thus, energy production is maintained, even when the oxidation of individual fuels is increased, implying reduced oxidation of competing fuels (competitive catabolism).

Next, we evaluated whether the competitive catabolism equation accurately quantifies the relationship between experimentally measured fluxes and concentrations. Using the parametrized competitive catabolism equation (Figure 1B and Tables S1-3), we predict catabolic fluxes on an individual mouse basis (across mice receiving different rates of ^13^C-nutrient infusions), based on the measured circulating metabolite concentrations. The predictions of the competitive catabolism equation closely matched the observed catabolic rates for 3HB, lactate, and glucose (Figure 2E, F; R^2^ = 0.93). This quantitative agreement further supports the validity of the competitive catabolism rate law *in vivo*.

### Detailed kinetic model

Competitive catabolism integrates the chemical principle of mass action with the biological one of energy homeostasis. Its implementation requires the concerted action of many dozens of metabolic enzymes. To explore how this might occur, we developed a differential equation model, operating at the level of individual enzymatic reactions, encompassing the catabolism of glucose, lactate, ketone bodies (3HB), and fatty acids (palmitate). The most comprehensive such model would include every reaction from genome-scale metabolic networks such as Recon3D,^29,30^ but dynamic models of this size (> 6,000 differential equations) are poorly tractable. We accordingly sought a simpler, unbiased alternative. To this end, we applied an algorithm to extract from Recon3D the minimal set of mass- and charge-balanced reactions required for the complete oxidation of each of the four above key metabolites^30–32^. The union of these reactions yields a mass-balanced network comprising 62 reactions, three compartments (extracellular, cytosol, mitochondria), and 70 metabolites.

To translate this network into a system of differential equations, we mathematically represented each reaction’s rate using Michaelis–Menten kinetics, incorporating textbook competitive inhibition and allosteric regulation^33^ (Figure 3A). Parameters were determined by sampling affinity constants (*K*_*m*_, *K*_*a*_, *K*_*i*_) randomly within a physiological range and calculating the corresponding maximal rate parameters (*V*_*max*_) to match experimentally observed concentrations and fluxes^34–36^. These parameter sets were then filtered to capture those that produce homeostatic models (consistent ATP production across varying nutrient levels and rapid return to a steady state following perturbation).

**Figure 3.**
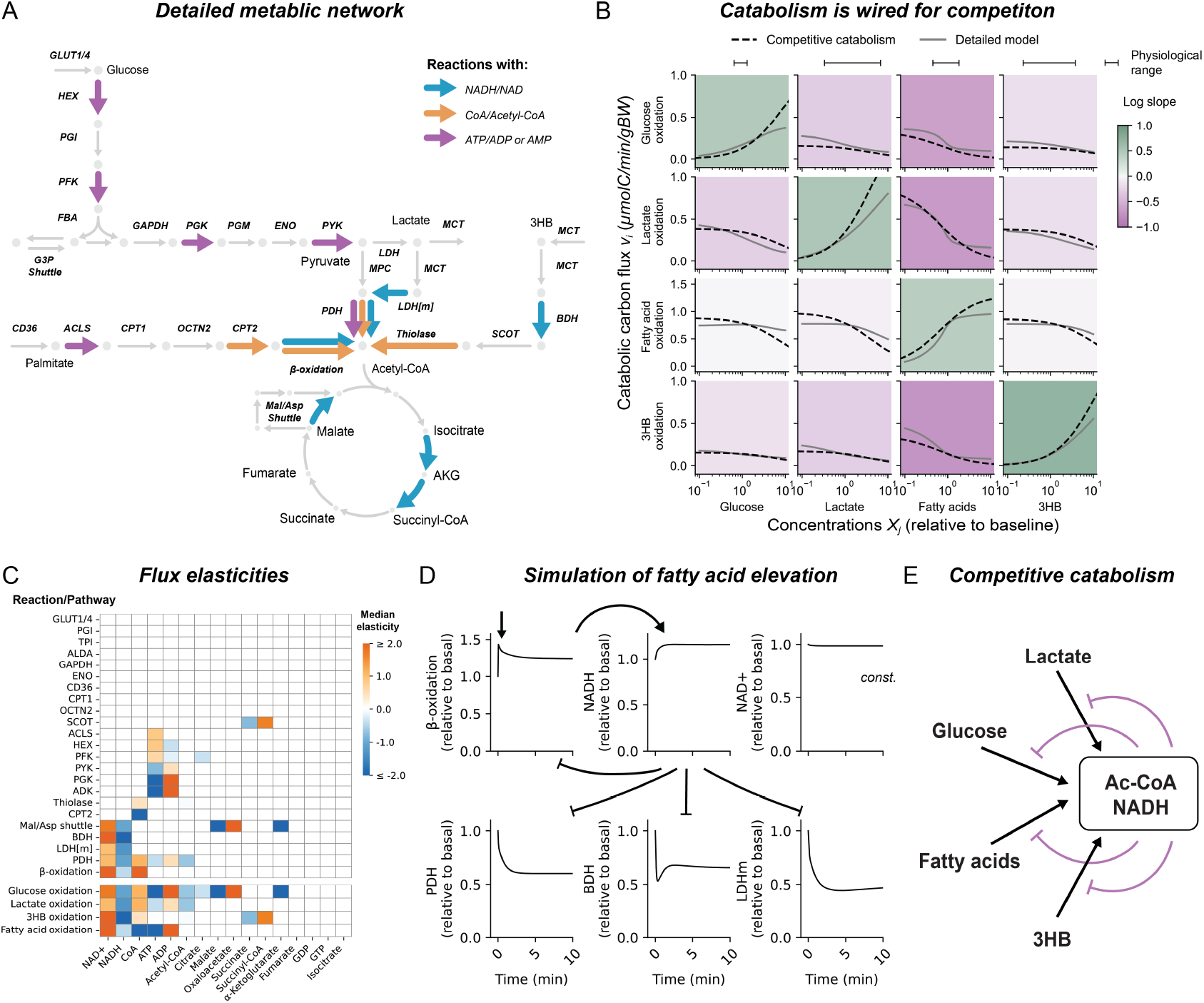
Detailed kinetic model of nutrient catabolism. A) Metabolic network that enables catabolism of glucose, lactate, ketone bodies, and fatty acids. Reactions involving shared cofactors are highlighted in color. B) Competitive catabolism is hardwired in the metabolic network architecture. Plots show the simulated impact on metabolic flux of changing the concentration of the metabolite shown on the X-axis (green boxes, mass action; purple boxes, competitive catabolism). Gray solid line represents the output of a characteristic detailed kinetic model (closely resembling the median behavior across sample models). Dashed black lines represent the competitive catabolism equation. C) The shared cofactor pairs NADH/NAD and AcCoA/CoA stand out for impacting the catabolic flux of all the examined nutrients (glucose, lactate, ketones and fatty acids). Heatmap shows the impact of metabolite levels on individual reaction fluxes calculated using the flux expressions of the detailed model (flux elasticities). The most strongly affected step in each nutrient’s catabolic pathway is highlighted in the bottom four rows. D) An increase in circulating fatty acids triggers elevated NADH and thereby inhibits catabolism of glucose, lactate, and 3HB. Plots are dynamical simulations based on the detailed kinetic model of an instantaneous 5-fold increase in fatty acids. E) Shared cofactors mediate competitive catabolism.

The above process yielded robust detailed kinetic models of nutrient catabolism. Importantly, these models were neither designed nor selected to show competitive catabolism. Nevertheless, they naturally show such behavior. Specifically, a rise in the concentration of any circulating nutrient both promotes that nutrient’s burning and suppresses other nutrients’ catabolism. Moreover, catabolic pathway fluxes calculated via the detailed kinetic model versus via the competitive catabolism equation agreed quantitatively (Figure 3B, and S3A).

### Mechanisms underlying competitive catabolism

We next sought to understand the mechanistic basis for competitive catabolism. First, we asked whether competitive catabolism depends on metabolic regulation, or purely the metabolic network’s biochemical connections. To this end, we removed all regulation from the detailed kinetic model and repeated the parameter identification process. Without regulation, far fewer parameter sets yielded robustly homeostatic models, highlighting the value of regulation for network robustness to parameter variation (Figure S3C,D). Nevertheless, some models lacking regulation were homeostatic, and such models manifested competitive catabolism (Figure S3B). Thus, competitive catabolism does not depend on metabolic regulation and instead is encoded in the architecture of the metabolic reaction network.

A logical way for pathways to compete, independent of regulation, is through shared substrates or products. Quantitative determination of the sensitivity of individual enzyme fluxes to changing metabolite levels (“elasticities”), identified only two pairs of metabolites that impact flux across all four catabolic pathways: the cofactors NADH/NAD and acetyl-CoA/CoA (Figure 3A). Both pairs impact nutrient catabolism upstream of the TCA cycle, allowing a properly balanced carbon flow into TCA oxidation. For both pairs, the low-energy form promotes catabolic flux, and the high energy form suppresses it (Figure 3C), with flux control residing in the less abundant (and hence more variable) member of the cofactor pair (e.g. NADH).

To illustrate the molecular events leading to competitive catabolism, we simulated a sudden elevation in fatty acid levels (Figure 3D). When fatty acids rise, β-oxidation increases, leading to higher NADH, whose accumulation leads to product inhibition of key dehydrogenases across all three competing pathways: pyruvate dehydrogenase (PDH), lactate dehydrogenase (LDH), and β-butyrate dehydrogenase (BDH). Additionally, elevated NADH transiently inhibits β-oxidation itself, until the system stabilizes at a new steady state characterized by increased fatty acid oxidation and suppressed catabolism of alternative fuels. The same basic response pattern holds for elevations in other nutrients (Figure S4): increased NADH (or, analogously, depleted CoA) suppresses alternative pathways (Figure 3E). Thus, competitive catabolism is mediated through shared cofactors

### Suppression of insulin action by nutrient excess

We next explored the role of competitive catabolism in insulin action. The gold-standard experimental approach for measuring insulin responsiveness is euglycemic-hyperinsulinemic clamp. In such experiments, insulin is steadily infused while glucose levels are maintained constant through manual adjustment of the rate of an exogenous glucose infusion by the experimenter^37,38^. The required glucose infusion rate serves as a quantitative measure of insulin responsiveness. To simulate this process computationally, we deployed our initial reductionist multi-nutrient dynamic model (Figure 1), modeling the experimentalist as a feedback controller that increases the glucose infusion rate when blood glucose falls below a setpoint of 100 mg/dL. Without any free parameters or fitting, the multi-nutrient model effectively mimicked the response of fatty acids (decreased), lactate (minimal change), and ketones (decreased) during the clamp, as well as the extent of required glucose infusion at two different insulin doses (1.25 mU·min^−1^·kg^−1^, which is about twice fasting insulin, and 2.5 mU·min^−1^·kg^−1^ which is typical postprandial insulin).

Insulin can induce glucose consumption either directly (by promoting glycolysis) or indirectly (by blocking lipolysis). A key role for lipolytic suppression is hinted at by the capacity for fatty acids to suppress insulin-stimulated glucose oxidation, first described by Randle in 1963^39^. While Randle originally proposed an allosteric mechanism for this effect (inhibition of phosphofructokinase by fatty acid-derived citrate), more recent studies favor signaling impairment (insulin signaling inhibition by intracellular lipids)^40–42^. We were curious if competitive catabolism could account for this effect. We replicated the classical experiment of infusing fat (intralipid) during euglycemic-hyperinsulinemic clamp. In parallel, this experiment was simulated using the multi-nutrient model with addition of an exogenous fatty acid influx. The simulation predicted that exogenous fatty acids would decrease the required glucose infusion to maintain euglycemia by about 75% at a low insulin dose, with a lesser effect at higher insulin dose, in good accordance with experimental results (Figure 4A,B and S5A,B).

**Figure 4.**
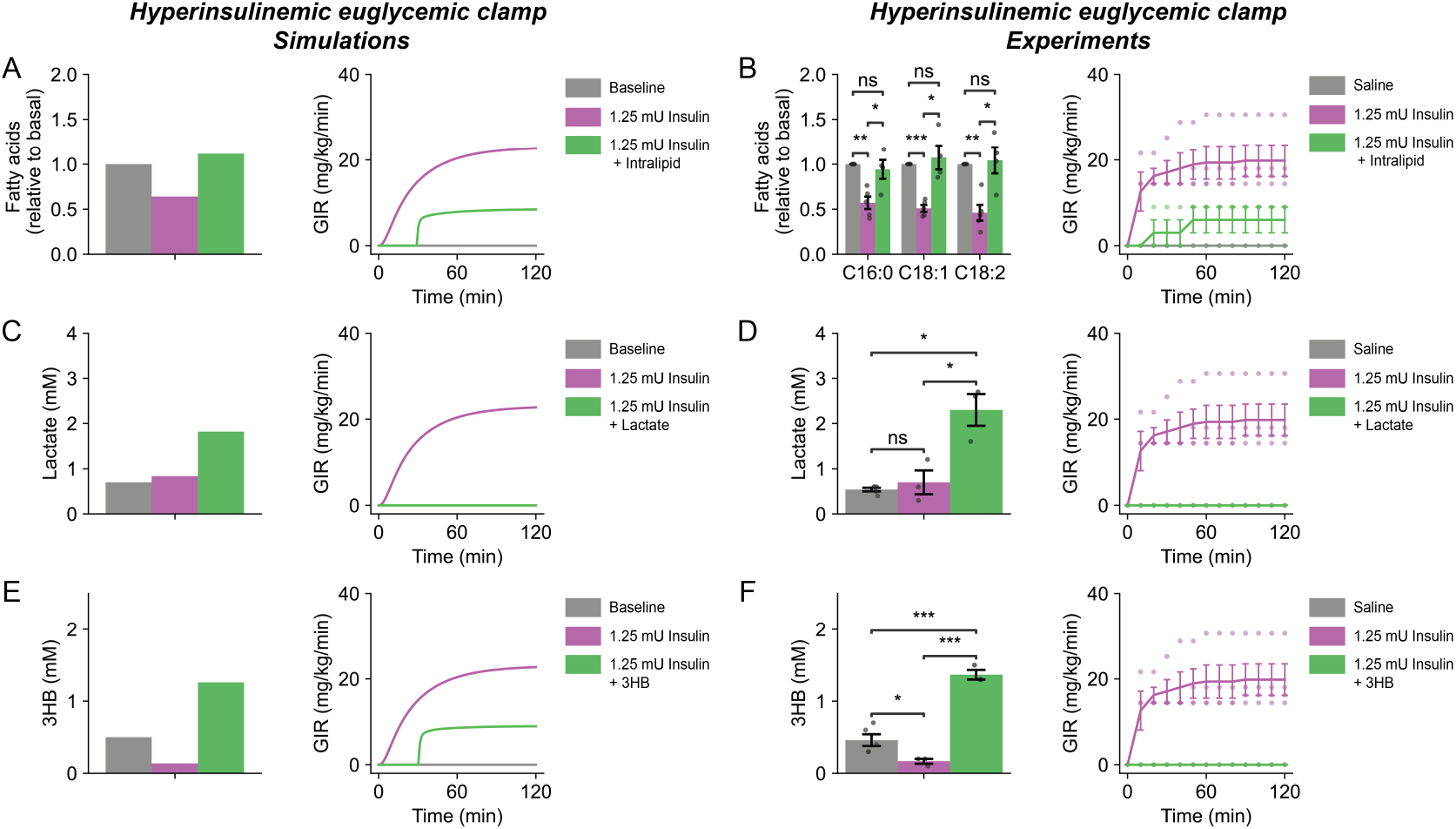
Acute insulin resistance induced by competitive catabolism. Simulated and experimental euglycemic-hyperinsulinemic clamps (steady insulin infusion with variable glucose infusion to maintain euglycemia). In addition to glucose infusion, perturbative intralipid, lactate, or 3HB infusion were also infused as indicated. The exogenous nutrient infusions trigger competitive catabolism and thereby suppress insulin-induced glucose consumption. Simulations are directly from the multi-nutrient model in Fig. 1 and involve no fitted parameters. The same data for saline (n = 5 mice) and insulin (n = 4 mice) are repeated in multiple panels. A) Simulation with intralipid infusion (simulated as fatty acid infusion). B) Experiment with intralipid infusion (n = 3 mice). C) Simulation with lactate infusion. D) Experiment with lactate infusion (n = 3 mice). E) Simulation with 3HB infusion. F) Experiment with 3HB infusion (n = 3 mice). 3HB, 3-hydroxybutyrate. ns, not significant. *p < 0.05, **p < 0.01, ***p < 0.001, by two-sample t-test. Data are mean ± SEM.

Intriguingly, the ability of alternative nutrients to suppress insulin-induced glucose uptake is not limited to fat. Lactate can also do so.^43–46^ This effect of lactate is hard to explain by lipid-mediated insulin signaling inhibition but aligns naturally with the concept of competitive catabolism. Indeed, in agreement with experimental results, under clamp conditions, the multi-nutrient model predicted that perturbative infusion of lactate (130 nmol min^−1^ [g body weight]^−1^, producing a rise in lactate to ~2.5 mM) would abolish the requirement for exogenous glucose to maintain euglycemia at the low insulin dose and mitigate it at the higher insulin dose (Figures 4C,D and S5C,D). Both predictions of lactate blood levels and required glucose infusion rates align well with experimental data. Thus, the multi-nutrient model has substantial predictive power.

The effects of fat and lactate on clamp responses were previously known. Accordingly, we sought a further prediction from the model: How would excess ketone bodies impact insulin sensitivity? Simulated perturbative ketone body infusion (120 nmol min^−1^ [g body weight]^−1^ of 3HB) was predicted to increase 3HB levels to about ~1.3 mM and to decrease the required glucose infusion by about 70% for the low insulin dose and 20% for the higher insulin dose. Experimental data conformed well to these predictions, showing complete ablation of glucose demand at the low insulin dose and strong suppression at the higher insulin dose (Figure 4E,F and S5E,F). Thus, simulations and experimental data demonstrate that, consistent with competitive catabolism, excess nutrient availability acutely triggers insulin resistance.

### Lipolysis suppression by insulin

A key feature of organismal metabolism is the ability to meet varying energy demands. Dynamic metabolic models have rarely, if ever, been tested for this capacity. We evaluated the ability of the multi-nutrient model to respond to an order-of-magnitude range of ATP demand, consistent with physiological variation in human energy expenditure (rest vs. physical activity)^47^. Consistent with exercise promoting glucose homeostasis, increased energy demand tended to lower glucose. The modeled variation in circulating glucose as a function of energy demand somewhat exceeded actual acute physiological variability, likely reflecting exercise adaptation also involving non-modeled factors (Figure 5A,B). Nevertheless, the homeostatic capacity of the multi-nutrient model was impressive.

**Figure 5:**
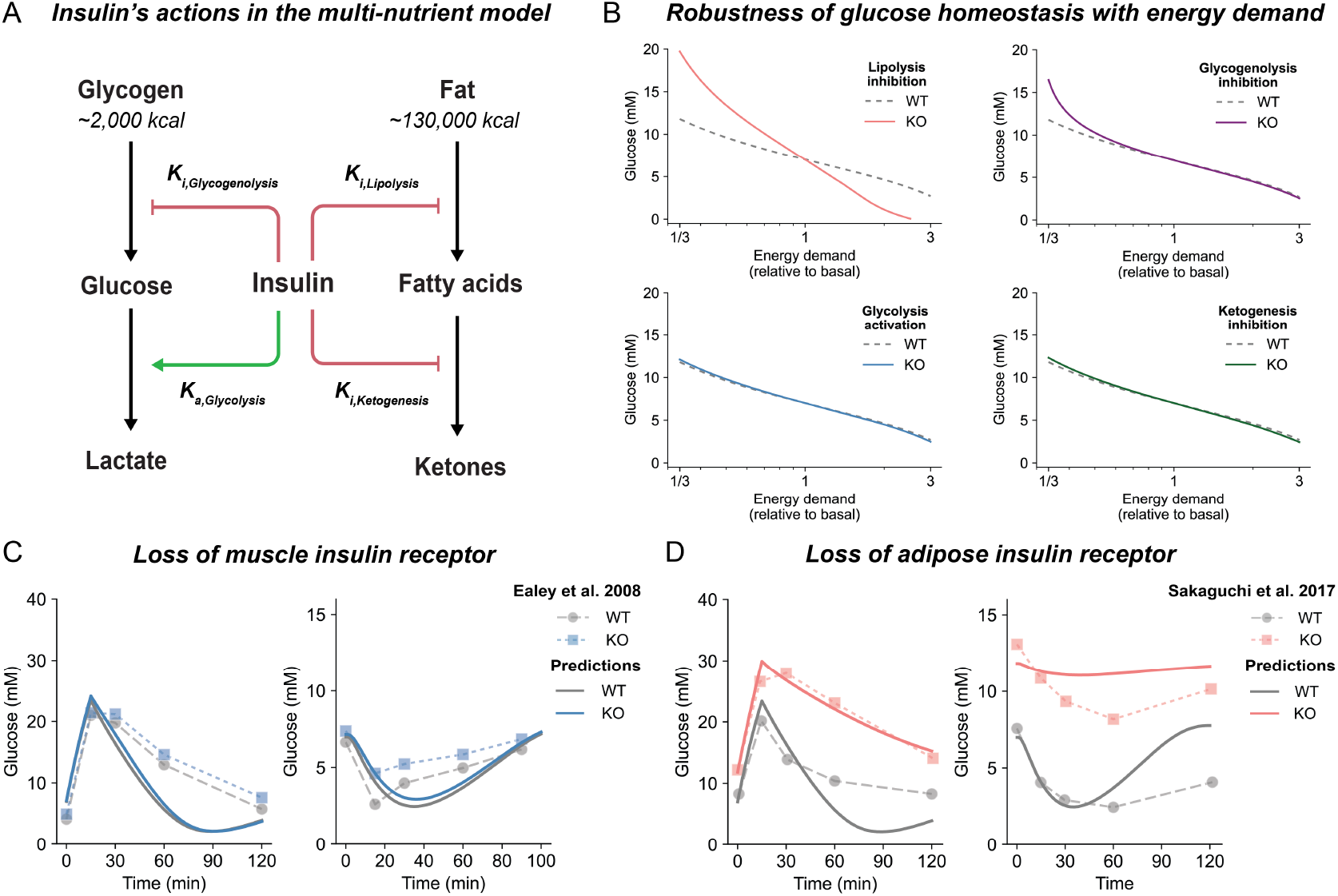
Dissecting insulin action on adipose versus muscle. A) Insulin has four distinct actions that are simulated quantitatively in the multi-nutrient model. B) In the multi-nutrient model, insulin action on adipose (lipolysis suppression) is required for robust glucose homeostasis. Plots show simulations of steady-state blood glucose as a function of whole-body energy demand, for the wild-type multi-nutrient model and variant models lacking each of insulin’s four actions. C) Loss of muscle insulin action has limited effect. Multi-nutrient model predictions and experimental data for glucose and insulin tolerance tests in mice with or without muscle insulin receptor (simulated as model without direct insulin activation of glycolysis)^89,90^. D) Loss of adipose insulin action causes insulin resistance. As in C for animals without adipose insulin receptor^91^ (simulated as model without insulin suppression of lipolysis).

This homeostatic capacity depends on insulin. We next explored which of insulin’s actions are required by individually removing each of insulin’s effects from the model. Across the tested range of energy demand, models lacking insulin-mediated glycolysis activation or ketogenesis suppression produced normal steady-state glucose levels and simulated glucose tolerance tests (GTT) and insulin tolerance tests (ITT) (Figure 5B,C). The ability of the multi-nutrient model, with no free parameters or data fitting, to mirror GTT and ITT results, is notable. Moreover, the robustness of the simulated GTT and ITT responses to loss of muscle insulin action aligns with murine experiments in which insulin receptor is genetically ablated selectively in muscle (Figure 5C) and with human genetic data showing that fasting glycemia is largely unaffected by mutations that impair insulin-induced GLUT4 translocation^48^ (though these mutations do slow return to euglycemia following carbohydrate feeding). Thus, the model captures the capacity for insulin to mediate fasting glucose homeostasis through mechanisms independent of direct glycolysis activation in muscle.

The effects of insulin on fasting glycemia could potentially occur instead through regulation of stored nutrient release. Consistent with this, models lacking insulin-mediated glycogenolysis suppression or lipolysis suppression showed altered fasting glucose homeostasis. The impact of glycogenolysis suppression was modest. Lipolysis suppression, in contrast, was crucial, with its loss resulting in a pathologically steep relationship between glucose levels and energy demand (Figure 5B): hyperglycemia when energy demand is low and severe hypoglycemia when demand rises. Loss of lipolysis suppression (i.e. adipose insulin signaling) also led to profound defects in simulated GTT and ITT. These quantitative predictions of the multi-nutrient model align with experiments in which the insulin receptor gene is selectively ablated in adipose (Figure 5D). Thus, insulin’s primary fasted effect is on lipolysis, with glucose controlled indirectly via competitive catabolism.

### A physiological circuit by which obesity causes hyperglycemia

Although the primary event in obesity is excessive fat storage, the strongest pathological linkage is to excess circulating glucose (diabetes). We were curious if the multi-nutrient competitive catabolism model – without invoking any biochemical defects such as impaired insulin signaling – would capture the obesity-hyperglycemia connection. To model the metabolic consequences of obesity, we assumed that triglyceride accumulation in adipose tissue leads by mass action to a proportional increase in the tissue’s propensity to lipolyze triglycerides (Figure 6A). This increased lipolytic propensity, modeled as the product of the mass action rate constant *k*_*lipolysis*_ and the body fat mass [*FM*], reflects a greater tendency to release fatty acids at any given level of insulin (i.e. an upward shift in the lipolysis-insulin curve, Figure 6A). The actual lipolytic flux is then determined by the quantitative interplay between metabolites and insulin encoded in the dynamic multi-nutrient model.

**Figure 6:**
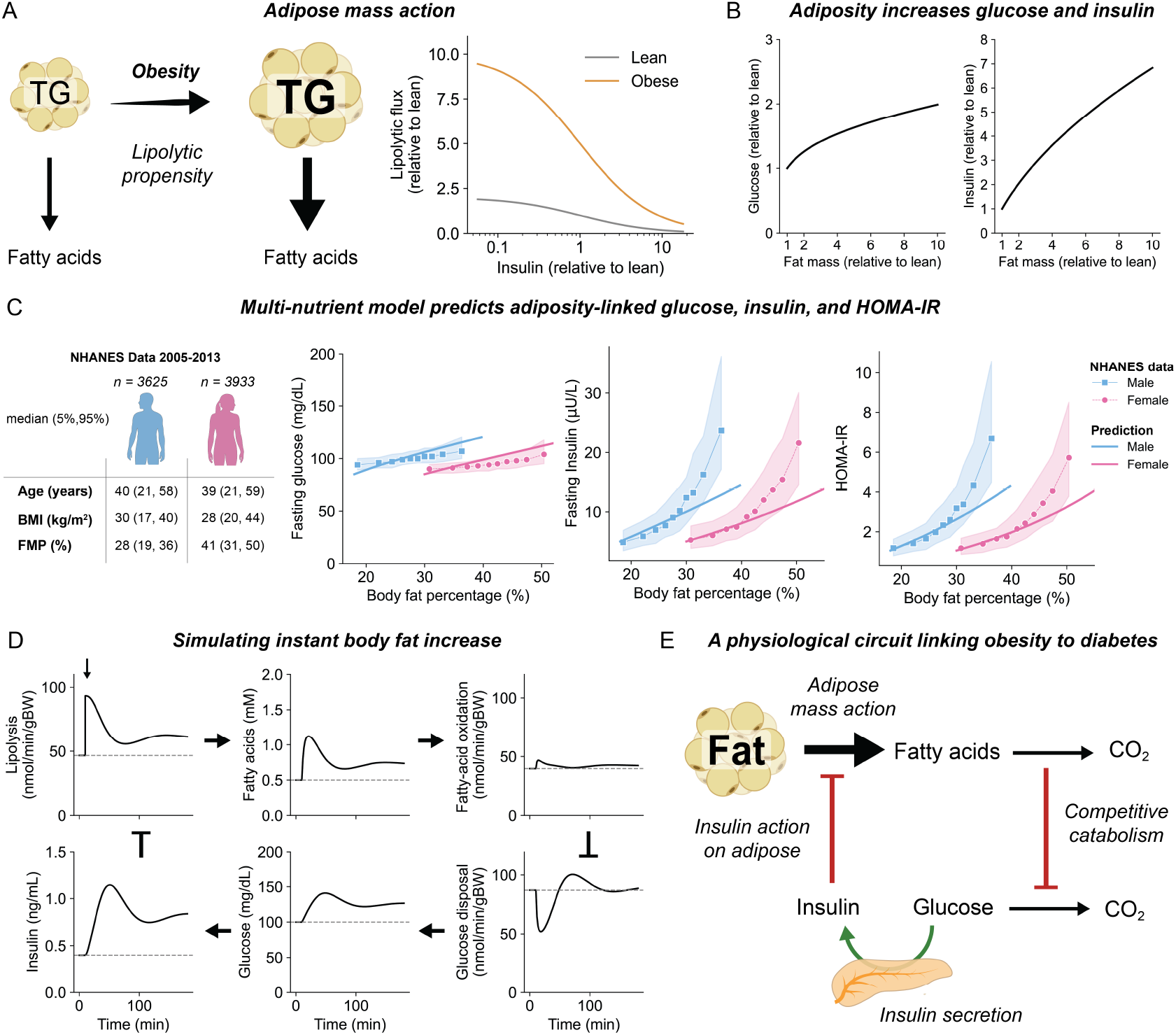
A physiological circuit linking obesity to insulin resistance. A) Due to mass action, increased fat mass enhances lipolytic propensity (lipolytic flux for a given level of insulin). B) The multi-nutrient model predicts that obesity leads to hyperglycemia and hyperinsulinemia. C) The model-predicted relationship between adiposity and fasted blood glucose, fasted insulin, and HOMA-IR match human data. Shown are the human population median and interquartile ranges for glucose, insulin and HOMA-IR at the indicated body fat percentage and predictions from the multi-nutrient model (NHANES, National Health and Nutrition Examination Survey). D) Increased body fat triggers a cascade of events leading to hyperglycemia and hyperinsulinemia. Plots are dynamical simulations, based on the multi-nutrient model, of an instantaneous 2-fold increase in body fat (which is physiologically infeasible but mechanistically informative). Initially, increased body fat causes enhanced lipolytic flux and circulating fatty acids. Via competitive catabolism, this leads to rising glucose, triggering insulin release which suppresses lipolysis. Higher insulin restores fatty acid homeostasis but hyperglycemia and hyperinsulinemia persist. E) A physiological circuit links obesity to diabetes.

To probe if there is a physiological connection between obesity and glucose elevation, we simulated steady-state fasting metabolite and insulin levels across a 10-fold range of body adipose mass. We observed a monotonic increasing relationship between adiposity and blood glucose that is robust to random perturbations of any model parameter values. Similarly, insulin, increased robustly with adiposity (Figure 6B and S6A). Notably, a 3-fold increase in adipose mass suffices to drive overt hyperglycemia and hyperinsulinemia, consistent with observations in genetically diverse murine cohorts (Figure S6B).

Insulin resistance in humans exhibits significant sex differences,^49^ with females maintaining metabolic health at higher body fat levels (Figure 6C). To reflect this difference, we scaled the multi-nutrient model to a sex-specific basal adiposity of 18% for healthy males and 30% for healthy females^50^. With this simple adjustment, the model captures the core relationship between body weight and fasting glucose, insulin and insulin resistance (HOMA-IR) in humans. No other changes were required to the multi-nutrient model to simulate human weight gain, even though the model was parameterized initially based on murine metabolite concentration and flux data, consistent with strong conservation of the core modeled circuitry.

The utility of the multi-nutrient model for revealing events leading to diabetes is illustrated by simulating an acute, two-fold increase in adipose tissue mass (Figure 6D). This increase in adipose tissue mass doubles lipolytic propensity and initially doubles lipolytic flux. Circulating fatty acid concentrations accordingly increase. The elevated fatty acid levels induce fatty acid oxidation, but due to finite ATP demand, the increase in fatty acid oxidation flux is relatively modest (25%). Glucose utilization is simultaneously suppressed through competitive catabolism. As a result, glucose disposal declines by about 40%, leading to a progressive rise in blood glucose. The ensuing hyperglycemia stimulates insulin secretion, with plasma insulin levels roughly doubling over the course of the simulation. Elevated insulin suppresses lipolysis, initiating a feedback loop that ultimately stabilizes the system at a new steady state characterized by modestly elevated lipolytic flux and fatty acid levels, a return to basal fatty acid oxidation flux, and elevated fasting glucose and insulin.

Thus, the model reveals a physiological circuit by which increased lipolytic propensity is feedback inhibited by insulin, at the expense of insulin and glucose elevation (Figure 6D). This circuit comprises four key components: (i) increase in lipolytic propensity by *adipose mass action*, (ii) inhibition of glucose disposal via *competitive catabolism*, (iii) glucose-induced increase in *insulin secretion*, and (iv) inhibition of lipolysis by *insulin action on adipose*. This physiological circuit maintains fat catabolic flux across a wide range of body composition. A physiological consequence of this circuit is that obesity drives both hyperglycemia and hyperinsulinemia.

### Modeling potential solutions

We finally asked whether the multi-nutrient model could be mined for means of ameliorating metabolic syndrome. To this end, we systematically changed each model parameter and asked if this change exacerbates or reverses obesity-induced hyperglycemia, hyperinsulinemia, or insulin resistance. Consistent with human genetics of type 2 diabetes, blood glucose levels were most sensitive to parameters governing insulin secretion – specifically, how much insulin the pancreas secretes in response to a given blood glucose level. These parameters, however, did not alter modeled insulin resistance. Instead, insulin resistance was primarily influenced by (i) energy demand, (ii) lipolysis, (iii) free fatty acid reesterification (Figure S6C). Increased lipolytic propensity (fat mass action) worsened insulin resistance, whereas elevated energy demand or reesterification improved it.

Based on these results, we simulated four interventions to counter obesity induced hyperglycemia and insulin resistance, (i) administration of exogenous insulin, (ii) exercise, (iii) loss of fat mass, and (iv) enhanced reesterification (Figure 7A). We observed that typical doses of exogenous insulin (0.2-0.5 U/kg) lower glucose but have minimal impact on insulin resistance, while exercise, fat loss, and enhanced reesterification can reverse metabolic syndrome systemically by reducing both glucose and insulin resistance (Figure 7B).

**Figure 7:**
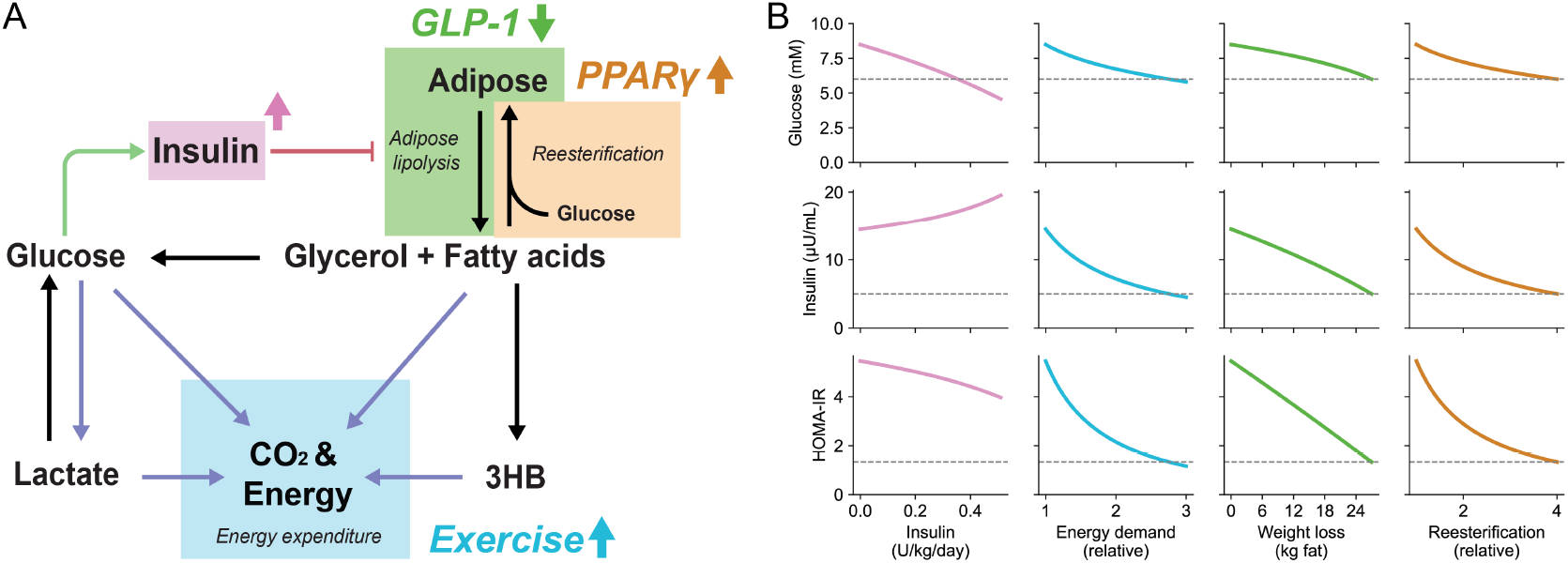
Modeling of potential solutions to resolve metabolic syndrome. A) Schematic of potential ways to ameliorate metabolic syndrome: purple = continuous insulin dosage, blue = increased energy expenditure; green = weight loss; orange = enhanced reesterification of circulating free fatty acids. B) Multi-nutrient model simulations for fasted glucose, fasted insulin and HOMA-IR, of approaches from (A). Dashed lines indicate corresponding values for a lean individual.

These model outputs align with the efficacy of weight loss interventions (e.g, via GLP-1 receptor agonists) and physical activity in reversing metabolic syndrome. Notably, increased energy expenditure has dual benefits: in addition to favoring weight loss, it directly promotes insulin sensitivity. The model further highlights reesterification as important to metabolic fitness. Reesterification removes free fatty acids from the bloodstream without generating ATP and thereby without triggering competitive catabolism. PPARγ agonist drugs (thiazolidinediones) stimulate adipocyte differentiation^51^, enhance whole body reesterification capacity, and (despite promoting weight gain) lower both blood glucose and lipid levels^52,53^. Thus, the model captures key axes for improving insulin sensitivity: appetite control, energy expenditure, and healthy adipose function.

## Discussion

Systemic metabolic homeostasis requires concerted regulation of the production and consumption of multiple nutrients. A key source of circulating metabolites is release from stored glycogen and fat.^28^ Insulin suppresses such release. It furthermore promotes glucose consumption. Consumption of many metabolites, however, appears substantially unregulated, mirroring chemical mass action.^23^ To examine how systemic metabolic homeostasis emerges without more extensive regulation of consumption fluxes, we developed a differential equation model, in which the overall activity of metabolic pathways is approximated by mass action (linear) functions of substrate concentration.

For catabolic pathways, this mass action approximation must be refined to account for total catabolic flux being set by energy expenditure, not metabolite concentrations. To achieve energy homeostasis, catabolism is a competitive process, wherein the burning of one nutrient suppresses that of others. The quantitative distribution of catabolic flux across pathways mirrors that of electric current across a parallel circuit of resistors. In such a circuit, each resistor carries a share of the total current, set by its conductance. Analogously, in catabolism, each pathway carries a share of the total flux, set by the product of the pathway’s catabolic propensity *k*_*i*_ and substrate concentration [*M*_*i*_]. Due to this concentration dependence, flux naturally adjusts to clear overabundant circulating nutrients. Our detailed kinetic model shows that cellular metabolism is hard wired for competitive catabolism. Thus, while different cell types have their own nutrient preferences (quantitatively reflected in the *k*_*i*_ values of the mass action approximation), our working model is that most cell types catabolize a wide variety of substrates in proportion to their availability. The net effect at the whole-body level is that catabolic fluxes self-regulate by preferentially oxidizing excess nutrients before ones that are scarce relative to their homeostatic set point.

An important advantage of modeling metabolism through mass action kinetics and competitive catabolism is that organismal metabolism is simulated with relatively few parameter values. Moreover, the required parameter values can be fully determined from a mixture of literature data regarding insulin action and *in vivo* metabolite concentration and flux measurements. This enabled the development of a predictive multi-nutrient model of systemic fasting metabolic homeostasis. This multi-nutrient model effectively simulates, without any free or fitted parameters, glucose and insulin tolerance tests; glucose clamps with and without additional perturbative nutrient infusions; tissue-specific insulin-receptor loss; and type 1 diabetes. Most informatively, the multi-nutrient model also predicts that obesity causes insulin resistance and type 2 diabetes. Extensive research has revealed numerous pathological mechanisms by which obesity impairs insulin action.^3–5^ The quantitative multi-nutrient model complements these pathological mechanisms with a physiological regulatory circuit by which obesity promotes insulin resistance without requiring defective insulin function.

This physiological circuit, centered around insulin, tailors lipolytic flux to energy demand. When lipolytic flux exceeds demand, circulating metabolite levels, including free fatty acids and glucose, rise. Glucose induces insulin which inhibits lipolysis. This resembles a classic negative feedback loop, except the feedback signal comes from glucose, even though the controlled pathway, lipolysis, makes free fatty acids. This dichotomy between pathway product (fatty acids) and feedback signal (glucose) is resolved by competitive catabolism, which links fatty acid and glucose concentrations. When fatty acids rise, glucose follows because its catabolism is competitively suppressed. Advantageously, this circuit design also leads to suppression of lipolysis immediately upon ingestion of a carbohydrate-containing meal, saving stored fat for future times of nutrient scarcity.

What happens in obesity? Increased triglyceride stores lead by mass action to a greater propensity for lipolysis. Cellular observations support a linear relationship between lipolytic flux and lipid droplet surface area^54^. The increased lipolytic flux and fatty acid levels indirectly elevate glucose and insulin. Elevated insulin in turn suppresses lipolysis, resulting in a metabolic steady state with only modestly elevated lipolytic flux and fatty acid levels but also hyperglycemia and hyperinsulinemia. Consistent with this model, in obesity, total fatty acid release is elevated, yet paradoxically suppressed when normalized to body fat mass.^55^ Moreover, circulating fatty acid levels are only modestly increased.^56^ Our quantitative model shows that this pattern is exactly what is expected based on the physiological regulatory circuitry. Thus, a fundamental cause of the obesity-diabetes relationship is the need for hyperinsulinemia to suppress the increased propensity for lipolysis resulting from excess stored fat.

While obesity is the most common cause of increased lipolytic propensity, defective fat storage can also lead to excessive lipolysis.^57–61^ Lipodystrophy is associated with limited fat stores, but pathologically elevated lipolytic flux and insulin resistance.^62^ Fat storage defects in lipodystrophy are particularly pronounced in the lower body.^63,64^ Effective lower body fat storage, observed clinically as a low waist-to-hip ratio, is associated high reesterification activity and metabolic health.^65–67^ While not directly encompassed in the current quantitative model, these observations further support a conceptual model where insulin sensitivity is substantially determined by the balance between reesterification capacity and lipolytic propensity.

### Limitations

The current multi-nutrient model is restricted to the fasted state and purely carbonaceous substrates. Modeling feeding requires accounting for nutrient storage processes including fatty acid, glycogen and protein synthesis that are currently omitted from the multi-nutrient model. Modeling nitrogen metabolism requires inclusion of glutamine synthesis and breakdown, transamination and the urea cycle. Adding these processes is an important future objective.

The current model is also generic, reflecting core metabolic reactions and regulatory processes that are conserved from mouse to human. Advantageously, and remarkably, this allows predictions of human phenotypes from the model parameterized using murine data, as the basics of competitive catabolism and insulin regulation are shared from rodents to humans.^68,69^ A long-term objective, however, is to deploy models of this sort to understand and predict individual patient phenotypes, including to facilitate the development of digital twins. This will require developing relationships between genetic polymorphisms and model parameter values and more fine-grained modeling of selected processes.

## Data and code availability

The authors declare that all the data supporting the findings of this study are available within the article and its supplementary information Files. Simulations can be reproduced and results accessed for academic use through an interactive web app (https://compcat.princeton.edu). Unprocessed experimental data underlying all plots and graphs are available in the supplemental data. The codes used to build and analyze the models are available on GitHub with adequate documentation (https://github.com/weilandtd/CompetitiveCatabolism).

## Author contributions

D.R.W. and J.D.R. conceived and designed the study. D.R.W. developed and analyzed the computational models, W.D.L., L.L., A.J.C. and C.H. performed mouse experiments, and D.R.W., W.D.L., and M.R.M. analyzed the data. Q.C. and W.D.L. performed mouse catheterization surgeries. J.A.B. provided access to indirect calorimetry and gas analyzers. N.S.W. contributed to theoretical analysis and data interpretation. D.R.W., and J.D.R. wrote the original draft. All authors discussed the results and commented on the manuscript.

## Materials and Methods

### Derivation of competitive catabolism

The competitive catabolism equation was derived to reconcile two paradoxical experimental observations – namely that nutrient oxidation increases with circulating concentrations (mass action), yet overall energy production remains constant. This derivation is grounded in a simplified model of catabolism that captures these two observations.

In cells, usable energy is primarily derived from the hydrolysis of ATP to ADP. Catabolism regenerates ATP from ADP such that the total adenine nucleotide pool is conserved, making ADP a shared resource for all catabolic pathways. This implies that catabolic flux in addition to the nutrient concentration depends on the availability of ADP. This relationship was expressed mathematically by considering that catabolism extracts energy by breaking down nutrients stepwise. In this model, each nutrient undergoes a series of intermediate reactions each contributing to ATP production. For illustrative purposes, this the simplest model assumes that each intermediate step yields one ATP. Thus, if catabolism of nutrient *M*_*i*_ yields *a*_*i*_ ATP, it is described by *n* = *a*_*i*_ sequential reactions:

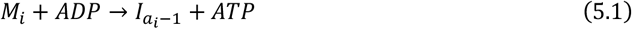

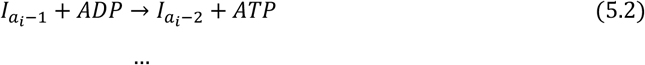

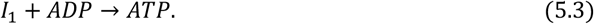

Catabolic intermediates within cells quickly settle into a new steady state after perturbation. It was therefore assumed that

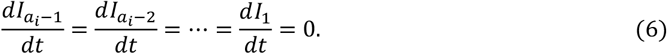

As a result, mass balance dictates that steady-state flux through each step is the same:

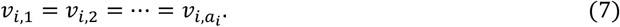

In the limit of irreversible reactions, flux through the pathway is then completely described by the reaction kinetics of the first step in the pathway *v*_*i*,1_. Assuming mass action kinetics the catabolic flux *v*_*i*_ of nutrient *M*_*i*_ is

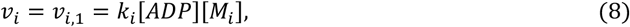

with the overall pathway stoichiometry

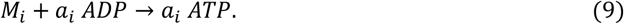

Simultaneously, ATP is constantly utilized at the rate of energy demand *v*_*energy*_. As mentioned above, the ATP produced from nutrient catabolism needs to meet this demand. Each nutrient *M*_*i*_ yield *a*_*i*_ ATP, thus, for *N* nutrients

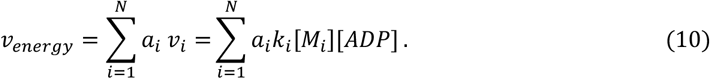

Because intracellular ADP turns over rapidly, it quickly settles into a new steady state, where

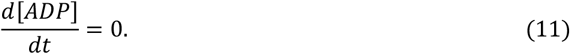

The steady-state concentration of ADP was then found by resolving equation (10):

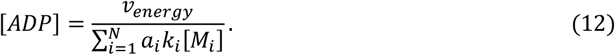

Combining equations (12) and (8), resulted in an expression for the catabolic fluxes, i.e., the competitive catabolism equation:

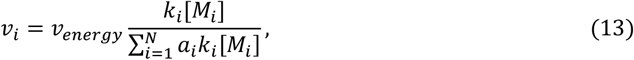

where *v*_*energy*_ is the whole-body energy demand, *k*_*i*_ are pathway-specific rate constants, [*M*_*i*_] are the circulating nutrient concentrations of the *N* nutrients, and *a*_*i*_ is the number of ATP derived from each nutrient. In this equation each catabolic flux *v*_*i*_ is bounded by its corresponding maximal catabolic rate *V*_*max,i*_ = *v*_*energy*_/*a*_*i*_. The flux *v*_*i*_ asymptotically approaches this limit when the respective nutrient concentration [*M*_*i*_] becomes large compared to its homeostatic setpoint:

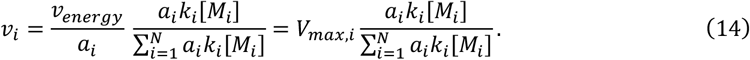

Catabolic fluxes are thus defined by their fractional contribution to the overall catabolic propensity 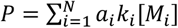. As a result, increasing the concentration of another nutrient [*M*_*j*_] decreases the catabolic flux *v*_*i*_ similarly to competitively inhibited Michaelis-Menten kinetics (Figure S1A).

### Building a model of fasted organismal metabolism

The model of fasted organismal metabolism was constructed by identifying the key production and consumption fluxes of the major circulating nutrients: glucose, lactate, fatty acids, and 3-hydroxybutyrate (3HB). Glucose and fatty acids are released through the breakdown of glycogen and triglycerides, respectively. Triglyceride catabolism also releases glycerol, which is continuously converted into glucose via gluconeogenesis. Conversely, reesterifying fatty acids into triglycerides requires glucose. Lactate is produced by tissues that preferentially rely on glycolysis and is subsequently either partially oxidized or recycled into glucose through gluconeogenesis. 3HB is synthesized from fatty acids via ketogenesis. All these nutrients can be oxidized to produce energy.

To relate the fluxes of these processes to the metabolite dynamics the above-described reaction network was translated into a set of differential equations for the circulating metabolite concentrations. Based on the principle of mass balance these equations relate the temporal rate of change in the molar amount of a species (i.e., number of molecules) to the molar fluxes (number of molecules per unit time) of its production and consumption (Eqn. 1). However, the dynamical variables are metabolite concentrations, expressed as molar amount per unit volume (e.g., mol/L), rather than absolute molecular counts. Consequently, the rate of change in the total number of molecules for a given metabolite is computed as the product of the rate of change of its concentration and the volume this amount is contained in, while the fluxes on the right-hand side are expressed in units of molar flux.

Importantly, circulating metabolites are not confined to the bloodstream but also bind to proteins and exchange with interstitial fluids and tissues. As a result, they are effectively distributed over a volume larger than the blood volume (i.e., volume of distribution^70^). Each metabolite has a distinct volume of distribution (*V*_*G*_, *V*_*F*_, *V*_*L*_, *V*_*K*_) which was determined experimentally (Figure S7, Table S2). These volumes were then used to express the differential equations in terms of the temporal rate of change in metabolite concentrations:

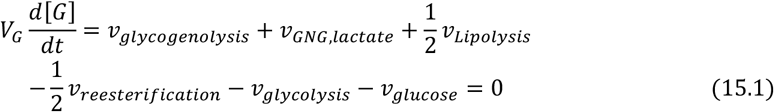

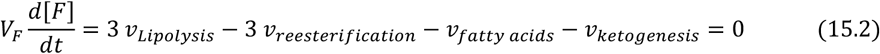

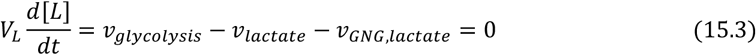

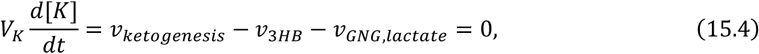

were [G] denotes glucose, [L] lactate, [F] fatty acids, [K] 3HB, concentrations in the circulation and *V*_*G*_, *V*_*L*_, *V*_*F*_, *V*_*K*_ are their respective volumes of distribution. Assuming the system operates at steady state, these mass balances constrain the reaction rates of the metabolic processes.

The energy producing fluxes *v*_*glycolysis*_, *v*_*glucose*_, *v*_*lactate*_, *v*_3*HB*_ and *v*_*fatty acids*_ are involved in competitive catabolism. Thus, their fluxes are constraint by the energy balance, i.e.:

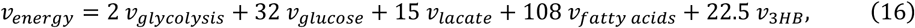

with numbers denoting the pathway specific ATP yields *a*_*i*_. All reaction fluxes were then determined by constraining the rate of reesterification to be two-thirds of the lipolysis rate^69^, and by matching the whole-body production rate of each nutrient to experimentally observed values^71^ (Table S3).

These mass balances are then converted into a system of ordinary differential equations by assigning mathematical expressions to the defined fluxes. Reaction rates were modeled using zero-order kinetics for fluxes originating from macromolecular reservoirs (*v*_*glycogenolysis*_, *v*_*Lipolysis*_), mass-action kinetics for anabolic and tissue-specific pathways (*v*_*GNG,lactate*_, *v*_*Reesterification*_, *v*_*ketogenesis*_), and competitive kinetics (Eqn. 14) for major catabolic routes involved in energy production (*v*_*glycolysis*_, *v*_*glucose*_, *v*_*lactate*_, *v*_*fatty acids*_, *v*_3*HB*_). Following the definition of chemical driving forces, insulin regulation was incorporated by multiplying each relevant flux with a regulatory term that, with increasing insulin, enhances glycolysis and suppresses glycogenolysis, ketogenesis, and lipolysis. Inhibition by Insulin (insulin action *I*_*a*_) is modeled as:

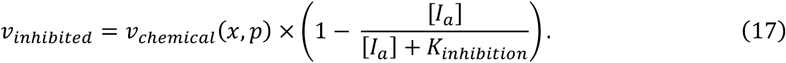

Insulin’s activation is modeled by a saturating activation term of the form:

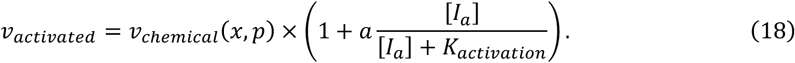

To maintain the energy balance constraint in the competitive catabolism equations, the glycolysis activation term must also be included in the denominator (Table S1).

Insulin dynamics were captured using two distinct variables: (i) circulating insulin [*I*], and (ii) insulin action [*I*_*a*_] allowing differentiation between rapid insulin dynamics in circulation and delayed peripheral signaling. Circulating insulin dynamics were modeled using Hill kinetics to capture glucose-dependent secretion and mass-action kinetics to represent degradation. Model parameters were chosen to reproduce experimentally observed insulin secretion and clearance dynamics:^23,72^

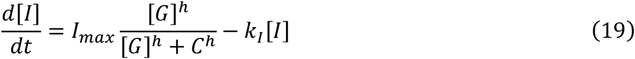

Insulin action follows circulating insulin with first-order delay (*τ* = 30 min):

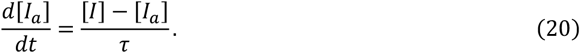

Final differential equations were derived by scaling each reaction flux in the mass balance equations to reflect metabolite concentration changes, achieved by dividing the mass fluxes by their respective volumes of distribution (Equations 15.1–4).

Parameters were constrained by equating the expression of the respective reaction rates with the steady-state flux profile. This results in ten independent constraints that allow us to express the mass action parameters (*V*_*glycogenolysis*_, *k*_*GNG,lactate*_, *k*_*reesterification*_, *k*_*lipolyis*_, *k*_*ketogenesis*_, *k*_*glycolysis*_, *k*_*glucose*_, *k*_*lactate*_, *k*_*fatty acids*_, *k*_3*HB*_) as a function of the steady-state levels *X*_0_ = [*G*_0_, *L*_0_, *F*_0_, *K*_0_, *I*_0_, *I*_*a*,0_], the steady-state mass fluxes, and the five regulatory parameters *K*_*i,lipolysis*_, *K*_*i,glycogenolysis*_, *K*_*i,ketogenesis*_, *K*_*i,lipolysis*_, *K*_*a,glycolysis*_ and *a* (Table S3). With insulin dynamics (*I*_*max*_, *h, C, k*_*i*_, *τ*), circulating concentrations and fluxes constrained to match experimentally observed values, this reduces the free parameter space to only five free parameters which are chosen with biological rational (Table S2).

### Basins of attraction

The stability of the multi-nutrient model steady state was probed by perturbing metabolite and insulin concentrations pairwise within a plane spanned by two variables. A region containing the relevant perturbed concentrations was defined, e.g., between 0 and 2 times the steady-state concentrations of glucose and fatty acids, and *n* = 24 equally spaced points along the circumference of the rectangle encompassing the region of interest were generated. Dynamical simulations were then performed with initial conditions being the respective values for the two concentrations combined with the steady state concentrations of the remaining variables.

### Hyperinsulinemic-euglycemic clamp simulations

During *in vivo* insulin clamps, the experimentalist manually adjusts the glucose infusion rate to maintain euglycemia in the presence of elevated insulin concentration. This was modeled *in silico* using an asymmetric p-controller, which infuses glucose proportionally to the deviation from normoglycemia (*G*_0_ = 100 mg/dL) when glucose levels fall. Mathematically the exogenous glucose infusion rate was expressed as:

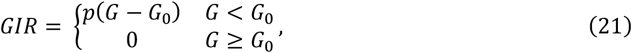

with *p* = *v*_*energy*_/*G*_0_ and a wide range of *p* given similar results.

### Simulation of glucose and insulin tolerance tests

Glucose tolerance tests were simulating by assuming complete absorption of intraperitoneally injected glucose over a 15-minute period at a constant rate. The absorption rate *v*_*IP*_ = 420 nmol min^-1^ (g body weight)^−1^ was chosen to match the initial rise in plasma glucose concentrations observed in the control animals. The simulation was performed by numerically integrating the system of differential equations for 15 minutes with constant glucose input, followed by an additional 105 minutes with the absorption rate set to zero.

Insulin tolerance tests were simulated by modeling the injection as an instantaneous increase in circulating insulin concentration to 60 ng/mL, which mirrors experimental data^13^, and simulating the dynamical response without further insulin input.

### Sensitivity analysis of the multi-nutrient model

Parameter sensitivities were calculated as scaled partial derivatives of metabolite concentrations or other observables with respect to individual model parameters:

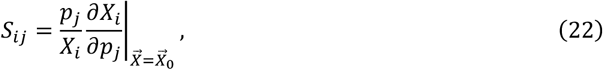

were *X*_*i*_ is the concentration or observable, *p*_*j*_ is model parameter, and 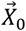 denotes the vector of steady-state concentrations. Partial derivatives were approximated using finite differences with a central difference scheme. To assess sensitivities under obese conditions, steady-state concentrations of metabolites and insulin were first computed at an elevated fat mass ([*FM*] = 3*x basal*). Sensitivities were then calculated based on this altered steady state.

### Minimal subnetwork extraction

In preparation for building a detailed kinetic model of catabolism, we first established a minimal reaction network sufficient for complete catabolism of the relevant nutrients. The lumpGEM algorithm was adopted to extract from a genome-scale metabolic model (GEM) the minimal set of thermodynamically feasible, charge- and mass-balanced reactions required for the complete oxidation of a metabolite. LumpGEM extends the thermodynamic flux analysis (TFA) framework to identify the smallest subnetworks that must be added to a core set of reactions to perform a biosynthetic task.^31^ TFA is a mixed-integer linear programming (MILP) formulation of flux balance analysis (FBA) that incorporates thermodynamic constraints on flux directionality using measured or estimated Gibbs free *energ*ies and metabolite concentrations.^73,74^ Here, lumpGEM was applied to identify the smallest reaction set capable of completely oxidizing an input metabolite (glucose, lactate, palmitate, or 3-hydroxybutyrate) within a thermodynamically curated version of Recon3.^29,30^ To ensure complete oxidation, the CO2 production flux was constrained to be equal to the product of the number of carbon atoms in the input metabolite and its input flux. Core reactions were defined as those comprising the electron transport chain and NADH-shuttling between cytosol and mitochondria (lactate shuttle, malate-aspartate shuttle, and glycerol-3-phosphate shuttle). The lumpGEM algorithm was then used to enumerate all reaction sets of minimal size for which complete oxidation of the respective metabolite is feasible. All minimal-size sets were then combined into a metabolite-specific superset. The union of these supersets and the core reactions was then used to build a TFA model that served as the basis for the detailed model.

### Thermodynamic flux analysis

We next set out to determine physiologically relevant compartmentalized fluxes and concentrations for every reaction in the detailed catabolic model constructed above. Thermodynamically constrained steady-state concentration and flux profiles were explored using the pytfa package.^75^ Metabolite concentrations were integrated into the model as natural logarithms of the concentrations:

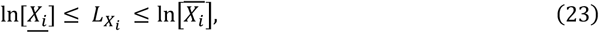

where 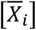 and 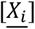 denote the upper and lower concentration bounds computed as mean ± standard deviation respectively, and 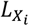 is a model variable representing the logarithmic concentration ln[*X*_*i*_]. Ratios of metabolite concentrations were constrained by formulating the equalities in logarithmic space, e.g.,

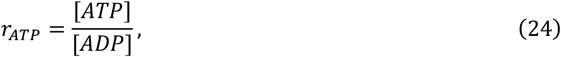

was constrained in terms of the logarithmic concentration variables *L*_*ATP*_ and *L*_*ADP*_ such that

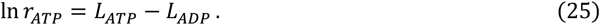

Reaction reversibilities were limited by constraining the concentration-dependent free energy of reactions to observed or reasonable ranges. In total, 49 metabolite concentrations, 8 concentration ratios, and 45 reaction free *energ*ies were compiled from the literature and used to constrain the TFA model (Supplemental Data, Table 1). These constraints reduced the variability of 77 compartmentalized concentrations to less than one order of magnitude and enabled full determination of all 70 fluxes (Supplemental Data, Tables 2 and 3).

To establish relevant upper and lower bounds for every flux and concentration based on the implemented constraints, 1000 constrained flux and concentration sets were generated starting from randomly selected thermodynamically feasible balanced flux sets by a Markov chain Monte Carlo simulation^76^. Upper and lower bounds were then determined as the 5% and 95% percentiles of the resulting distributions.

### Building a detailed dynamic model of cellular catabolism

The TFA model was converted into a kinetic model using the SKiMpy software.^77^ Reaction rates were expressed as functions of metabolite concentrations based on the following principles: (i) Transport reactions of small molecules such as oxygen, and CO_2_ are modeled using reversible mass-action kinetics. (ii) All other reactions were encoded using generalized saturable rate equations (e.g. Michaelis-Menten) whenever possible using reversible hill kinetics^78^ without cooperativity (Hill coefficient = 1). For reactions involving more than two substrates and products, or an unequal number of substrates and products, convenience kinetics^79^ were used. Subsequently, canonical enzyme-metabolite interactions (Supplemental Data, Table 4) were integrated by multiplying the chemically defined rate expression with regulatory terms, as described above^77^. The resulting kinetic model comprised 52 independent differential equations (68 mass balances with 16 conserved metabolite pools), involving 63 concentration-dependent reactions rates. The total concentration of the conserved pools is a conserved quantity that remains constant over time despite changes in individual metabolite concentrations (Supplemental Data, Table 5). Common examples include metabolites involved in energy (ATP, ADP, and AMP) and redox metabolism (NAD and NADH).^80,81^

To capture mitochondrial *energ*etics, one additional differential equation describing the dynamics of the mitochondrial membrane potential (MMP) was added. Considering the mitochondrial membrane as a capacitor, MMP dynamics are described fully by the net electric current across the membrane *J* and the membrane capacitance *C*. The electric current flows across the membrane in the form of ions. In mitochondria the predominant ions moving across the membrane are protons. Thus, the differential equation governing the MMP dynamics can be expressed in terms of all reactions that transport protons and charge across the membrane:

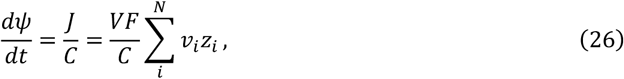

where *V* is the cell volume, *F* is Faraday’s constant, *C* is the membrane capacitance, *v*_*i*_ are reaction rates (mmol L^-1^ min^-1^), *z*_*i*_ are the number of elementary charges reaction *i* transports across the membrane. Signs of *z*_*i*_ were defined so that *ψ* is positive at steady-state and positively increases when positive charges are transported out of the mitochondria. The membrane capacitance was chosen so that the time constant of the equivalent resistance-capacitance circuit is 2 min.

The MMP dynamics were coupled to the electron transport chain reactions by expressing their effective thermodynamic driving force as a function of the MMP. This was achieved by modifying the apparent equilibrium constant of each charge-transporting reaction to depend on both the mitochondrial membrane potential and the net charge transported across the membrane:

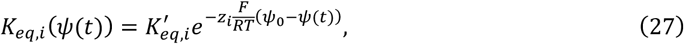

where 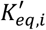 is the equilibrium constant computed from effective standard Gibbs free energy at the steady-state membrane potential *ψ*_0_, *ψ*(*t*) is the dynamic membrane potential, *z*_*i*_ is the reaction’s net charge transport across the membrane, *F* is the faraday constant, *R* the general gas constant and *T* the absolute temperature.

The resulting model was then parameterized using the standard Gibbs free *energ*ies from the TFA model, and by sampling the 313 affinity constants (*K*_*m*_, *K*_*a*_, *K*_*i*_). Parameters were sampled by drawing 20,000 independent random number vectors that were scaled to represent affinity constants (*K*_*m*_, *K*_*a*_, *K*_*i*_), and then back calculating the respective rate proportional parameters (*k, V*_*max*_) as previously described^34–36^. This resulted in 20,000 parameter sets that are consistent with the steady-state concentration and flux profiles. These parameter sets were then pruned based on local stability and linear dynamics: Unstable parameter sets, and parameter sets that had a larger than 20-minute characteristic timescale (largest eigenvalue of the Jacobian^34^) were rejected. The remaining parameter sets were then challenged by varying each nutrient concentration across a 100-fold range (10-fold up and down) and solving the differential equations to obtain the dynamic response. Time-dependent metabolite concentrations were computed by adjusting the concentration parameter and integrating the ODE system for 500 minutes (see below). Only parameter sets that reached a steady state and maintained a constant ATP production flux across all perturbations were retained for the final catabolic dose-response analysis (Figures 3 and S3).

### Dynamic analysis of detailed models

The detailed metabolic models were analyzed using the SKiMpy software.^77^ Concentrations were scaled units of mM and reaction rates to units of mM min^-1^. Time integration was performed using an implicit differentiation schema (Backward differentiation formula, BDF) with adaptive time step control as implemented in the CVODES solver library.^82,83^ Time steps are chosen dynamically by the algorithm to maintain a relative integration tolerance below 10^-6^. The final concentrations were considered to be a steady state if the absolute time derivatives 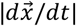 became sufficiently small (< 10^-2^ mM min^-1^) during the last integration steps.

To represent the characteristic behavior of the pruned parameter population, a representative parameter set was selected based on its similarity to the typical model output. Specifically, the coefficient of determination (R^2^) was computed between the median output of the population and each individual model output across all concentration-flux responses from the dose-response analysis (Figure S3A). The parameter set with the highest overall similarity (R^2^ = 0.997) was selected as the characteristic model.

### Animals

Animal studies followed protocols approved by the Princeton University Institutional Animal Care and Use Committee. All experiments were performed on 10–14-week-old C57BL/6N male mice (Charles River Laboratories). Aseptic surgery was performed to place catheters in the right jugular vein and left carotid artery connected to a vascular access button implanted under the skin on the back of the mouse (Instech Laboratories). Mice were allowed to recover from catheterization surgery for at least 5 days before experimentation. Mice with catheters were individually housed in environmentally enriched cages (The Andersons, Bed-r’Nest) with ad libitum access to water and food. Catheters were flushed with sterile saline and refilled with sterile heparin glycerol locking solution (SAI Infusion Technologies, HGS) every 5-6 days. Animals were housed on a normal light cycle (8AM-8PM) and fed a standard rodent chow (LabDiet, PicoLab Rodent 20 5053) provided on the floor.

### Plasma and tissue sampling

On the day of the experiment, mice were transferred to new cages with enrichment (The Andersons, Bed-r’Nest) and water gel (ClearH2O, HydroGel) but without food around 8AM (beginning of the light cycle). To minimize excess noise, vibrations, and other disturbances, mice cages were transferred to a quiet and isolated procedure room. Catheters were flushed with sterile saline. Experimenter left the room and let the mice acclimate to the new environment. Around 1PM, catheters were flushed again with sterile saline, and mice were weighed and connected to infusion lines with swivel and tether (Instech Laboratories products: swivel SMCLA, line KVABM1T/25) that allowed the animals to have free movement in the cage. Arterial and venous infusion lines were pre-filled with heparinized saline (10 U mL^−1^) and saline respectively before connecting to the catheters. Experimenter left the room, and mice were left in cages connected to lines for one hour to reduce stress. Around 2PM, a pre-determined dead volume (70 μL) was withdrawn from the arterial infusion line and arterial blood (10 μL) was sampled from the arterial catheter using a capillary blood collection tube coated with lithium heparin (Sarstedt, 41.1503.105). The arterial infusion line was refilled with the withdrawn dead volume (70 μL). Blood samples were stored on ice and then centrifuged at 10,000 × g for 10 min at 4°C to get plasma samples. Plasma samples were kept at −80 °C until LC-MS analysis.

For tissue samples, mice were euthanized by injecting 120 mg kg^−1^ pentobarbital via arterial catheter. All blood sampling was done before pentobarbital injection. Sodium pentobarbital 390 mg/mL (Vortech Pharmaceuticals, Ltd., FATAL-PLUS SOLUTION) was diluted in sterile saline to obtain a 120 mg/mL working solution. For a 25 g mouse, 25 μL of 120 mg/mL pentobarbital working solution was injected through the arterial catheter. The original FATAL-PLUS SOLUTION contains 29% v/v ethanol and 1% v/v propylene glycol. In the working solution, ethanol and propylene glycol concentrations are 8.9% v/v and 0.31% v/v, respectively. Tissues were quickly dissected and snap frozen in liquid nitrogen with pre-cooled Wollenberger clamp. Tissue samples were kept at −80 °C until LC-MS analysis.

### Plasma metabolite extraction

Plasma (2.5 μL) was added to 60 μL of −20°C 25:25:10 (v/v/v) acetonitrile:methanol:water solution, vortexed for 10 s, and put on ice for at least 5 min. The resulting extract was centrifuged at 21,000 × g for 20 min at 4°C and supernatant was transferred to tubes for LC-MS analysis. A procedure blank was generated identically without plasma, which was used later to remove background ions.

### Tissue metabolite extraction

Snap-frozen tissues were transferred to 2 mL round-bottom Eppendorf Safe-Lock tubes on dry ice. Samples were then ground into powder with a Cryomill machine (Retsch) for 1 minute at 25 Hz, maintained at a cold temperature using liquid nitrogen. For every 20 mg tissue, 800 μL of −20 °C 40:40:20 (v/v/v) acetonitrile:methanol:water with 0.1 M formic acid was added to the tube, vortexed for 10 s, and allowed to sit on ice for 10 minutes. For each 800 μL of tissue extract, 70 μL of ice-cold 15% (w/v in H_2_O) NH_4_HCO_3_ was added and vortexed to neutralize the sample. Samples were then centrifuged at 21,000 × g for 20 min at 4 °C. The supernatants were then transferred to plastic vials for LC-MS analysis. A procedure blank was generated identically without tissue and was used later to remove the background ions.

### Metabolite measurement by LC-MS

Metabolites were analyzed using a Vanquish Horizon UHPLC System (Thermo Scientific) coupled to an Orbitrap Exploris 480 Mass Spectrometer (Thermo Scientific). Waters XBridge BEH Amide XP Column (particle size, 2.5 μm; 150 mm (length) × 2.1 mm (i.d.)) was used for hydrophilic interaction chromatography (HILIC) separation. Column temperature was 25 °C. Mobile phases A = 20 mM ammonium acetate and 22.5 mM ammonium hydroxide in 95:5 (v/v) water:acetonitrile (pH 9.45) and B = 100% acetonitrile were used for both ESI positive and negative modes. The linear gradient eluted from 90% B (0.0–2.0 min), 90% B to 75% B (2.0–3.0 min), 75% B (3.0–7.0 min), 75% B to 70% B (7.0–8.0 min), 70% B (8.0–9.0 min), 70% B to 50% B (9.0–10.0 min), 50% B (10.0–12.0 min), 50% B to 25% B (12.0–13.0 min), 25% B (13.0–14.0 min), 25% B to 0.5% B (14.0–16.0 min), 0.5% B (16.0–20.5 min), 90% B (20.5 – 25.0 min). The flow rate was 0.15 mL/min. The sample injection volume was 5 μL. ESI source parameters were as follows: spray voltage, 3200 V or −2800 V, in positive or negative modes, respectively; sheath gas, 35 arb (arbitrary units); aux gas, 10 arb; sweep gas, 0.5 arb; ion transfer tube temperature, 300 °C; vaporizer temperature, 35 °C. LC–MS data acquisition was operated under full scan polarity switching mode for all samples. The full scan was set as: orbitrap resolution, 120,000 at m/z 200; AGC target, 1e7; maximum injection time, 200 ms; scan range, 60–1000 m/z.

### Non-perturbative and perturbative stable isotope tracer infusion

Stable isotope tracers (Cambridge Isotope Laboratories) were infused for 2.5 h through the implanted right jugular vein catheter. For minimally perturbative experiments, 2.5 (w/w in H_2_O) sodium-[U-13C]L-lactate, 130 mM [U-13C]glucose, or 100 mM [U-13C]3-hydroxybutyrate was infused at 0.1 μL min^−1^ (g body weight)^−1^. For intentionally perturbative measurements, 5% (w/w in H_2_O) sodium-[U-13C]L-lactate, 670 mM [U-13C]glucose, or 400 mM [U-13C]3-hydroxybutyrate was infused at 0.3 μL min^−1^ (g body weight)^−1^. The same fasting and sampling procedure was performed as described above.

### Indirect calorimetry and ^13^CO_2_ measurement

Indirect calorimetry was performed in individually housed pre-catheterized mice using a 20-channel open-circuit indirect calorimeter, in which 16 cages were mounted inside two thermally controlled cabinets, maintained at 22°C (Sable Systems International, Promethion). After overnight acclimation (20 h), oxygen consumption (VO_2_) and carbon dioxide production (VCO_2_) were measured during a 2 h saline or insulin infusion (clamp) after brief fasting (7AM-noon). For ^13^CO_2_ measurements, exhaled ^13^CO_2_ from minimal and perturbatively infused stable isotope tracers ([U-13C]glucose, [U-13C]L-lactate, and [U-13C]3-hydroxybutyrate) was measured via a stable isotope gas analyzer and arterial blood was sampled for measuring blood tracer labeling enrichment. The respiratory exchange ratio (RER) was calculated as the ratio of VCO_2_:VO_2_. Energy expenditure was calculated as heat (kcal h^−1^) = (3.815 + 1.232 x RER) x VO_2_). The metabolic studies were performed at the Penn’s Rodent Metabolic Phenotyping Core (University of Pennsylvania).

### Hyperinsulinemic-euglycemic clamp experiments

Hyperinsulinemic-euglycemic clamps were performed according to standard procedure^37,38^ with modifications. On the day of the experiment, pre-catheterized mice were fasted (5-6 hours) and connected to infusion lines with swivel and tether as described above. Around 2 PM, the insulin clamp began by infusing insulin or saline (control) and variable 1 M glucose via an implanted catheter in the right jugular vein. Insulin infusion rate was 1.25 mU min^-1^ (kg body weight)^-1^, unless otherwise stated. The insulin used, Humulin R (Eli Lilly, U-100), was diluted with sterile 0.1% BSA-saline solution to achieve concentrations of 12.5 mU mL^-1^, and the pump rate was set at 0.1 μL min^−1^ (g body weight)^−1^. Glucose solution (1 M) was prepared in sterile saline. Euglycemia was maintained (120-130 mg dL^-1^) by measuring arterial blood glucose (<5 μL) every 10 minutes with a glucose meter (Roche, Accu-Chek Aviva) and adjusting the glucose infusion rate as necessary. The insulin clamp was carried out for 2 h and blood samples (40 μL each) were collected at 110 and 120 minutes for downstream analyses.

For the insulin clamp with intralipid, 20% Intralipid (Sigma, I141) was infused at 0.01 μL min^−1^ (g body weight)^−1^ at the beginning of the insulin clamp, and variable 1 M [U-13C]glucose was infused to maintain euglycemia. The Intralipid (Sigma, I141) is composed of 18.0-22.0% wt soybean oil, 1.9-2.5% glycerin, 0.9-1.3% phospholipid, and ≤ 5.0 MEQ/L free fatty acids, with a pH range of 8.0-8.9. The soybean oil component is primarily composed of triglycerides, with a typical fatty acid composition of approximately 54% linoleic acid, 25% oleic acid, 11% palmitic acid, and 7% stearic acid.

### LC-MS data analysis

LC-MS raw data files (.raw) were converted to mzXML format using ProteoWizard (version 3.0.20315).^84^ El-MAVEN (version 0.12.0)^85^ was used to generate a peak table containing m/z, retention time, and intensity for the peaks. For tracer experiments, isotope labeling was corrected for ^13^C natural abundances using AccuCor2 package^86^. Statistical analyses were performed using the python statsmodels package.

### Endogenous production and disposal fluxes

To measure the endogenous production (R_a_) and disposal fluxes (R_d_) of a nutrient with a carbon number of *C*, the uniformly ^13^C-labeled form of the nutrient was infused. At steady state, the fraction of the labeled form [*M* + *i*] of the nutrient in plasma was measured as *L*_[*M*+*i*]_. The whole-body disposal flux (*R*_*d*_) is defined as

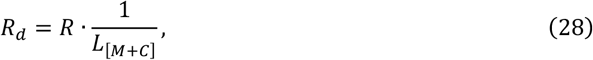

where *R* is the infusion rate of the labeled tracer. The whole-body production flux (*R*_*a*_) is defined as

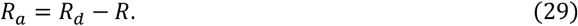

### Nutrient contribution to exhaled CO_2_

The stable isotope gas analyzer connected to the indirect calorimetry (Promethion, Sable Systems International, NV) reports the ratio of the heavy to light isotope (i.e., ^13^C/^12^C) in the breath sample in terms of δ^13^C (with units ‰), which is defined as:

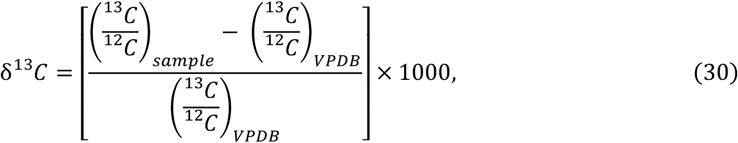

where (^13^C/^12^C)_VPDB_ is 0.0112372. VPDB (Vienna PeeDee Belemnite) is the international reference standard for carbon isotopes.^87^

For higher concentrations of ^13^C (e.g., above 100‰), fractional enrichment of ^13^C is usually expressed in terms of the ^13^C fraction (*L*),^88^ which is defined as:

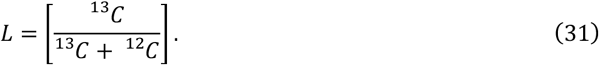

First, we converted the reported δ^13^C values to 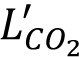 with the following equation:

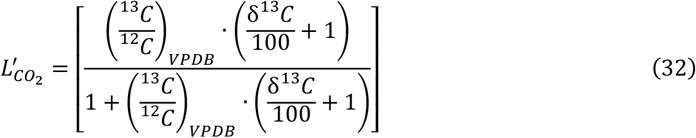

where (^13^C/^12^C)_VPDB_ is 0.0112372 as mentioned above. This value contains a background fraction from non-tracer sources (e.g., natural abundance), which is subtracted to compute the true CO_2_ labeling fraction from the infused tracer 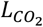:

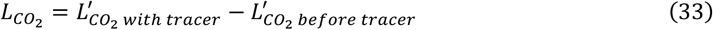

Next the total oxidation flux of the infused nutrient was calculated by normalizing the fraction of exhaled labeled 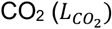 by the arterial^13^C enrichment of the nutrient (*L*_*n*_), and multiplying the result by the total rate of exhaled CO_2_:

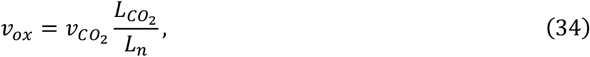

with *L*_*n*_ as

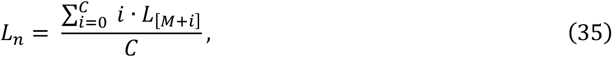

where C is the carbon number of the respective nutrient (e.g. C=3 lactate) and L_[M+i]_ is the fraction of the labeled form [M+i]. Note that there are C+1 labeled forms, such that *i* ranges from 0 to C.

### Volume of distribution of metabolites

Volume of distribution was measured based on circulatory flux (greater flux → bigger volume), circulating concentration (higher concentration → smaller volume), and the half-time for circulatory labeling to reach steady state during continuous tracer infusion (longer turnover time *τ* → bigger whole body metabolite pool → larger volume). To determine the turnover time *τ*, after a 5-6 h fast, [U-^13^C]-labeled tracers were infused to double catheterized mice at 3 μL min^-1^ for 6 minutes to fill the jugular vein catheter and remove the dead volume (~13 μL). The mice were then left in cages with the infusion lines attached for another 1.5 h to clear any residual tracer in the bloodstream. Thereafter the continuous tracer infusion was initiated and arterial blood sampled serially to determine in the labeling kinetics. The steady state labeling (*L*_∞_) and the turnover time (*τ*) were determined by fitting an exponential decay to the time dependent labeling:

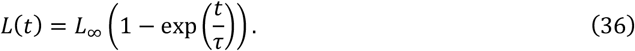

From the steady state labeling *L*_∞_ the metabolite circulatory flux *F*_*circ*_ can be determined from the infusion rate of the metabolite:

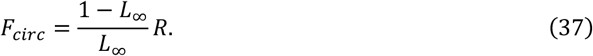

The respective whole-body metabolite pool size *P* can be determined as:

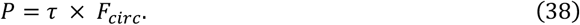

Distribution volumes of glucose, lactate, and fatty acids were calculated as:

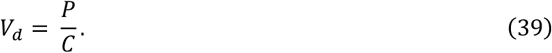

where *V*_*d*_ is the volume of distribution and *C* is the metabolite concentration in the arterial blood.

### Additional statistical analyses

Unless otherwise noted, data were analyzed using python (Packages: scipy, pandas, matplotlib, statsmodels, and seaborn) and are presented as mean ± standard error. P-values were calculated using unpaired two-tailed t-tests, with significance defined as \**p* < 0.05, \*\**p* < 0.01, \*\*\**p* < 0.001, and \*\*\*\**p* < 0.0001.

## Notes

https://github.com/weilandtd/CompetitiveCatabolism

https://compcat.princeton.edu

### Competing Interest Statement

The authors have declared no competing interest.

### Summary of Updates

To reflect the title in the pdf

