## Supplemental Figures and Tables for "Competitive catabolism in systemic metabolic homeostasis"

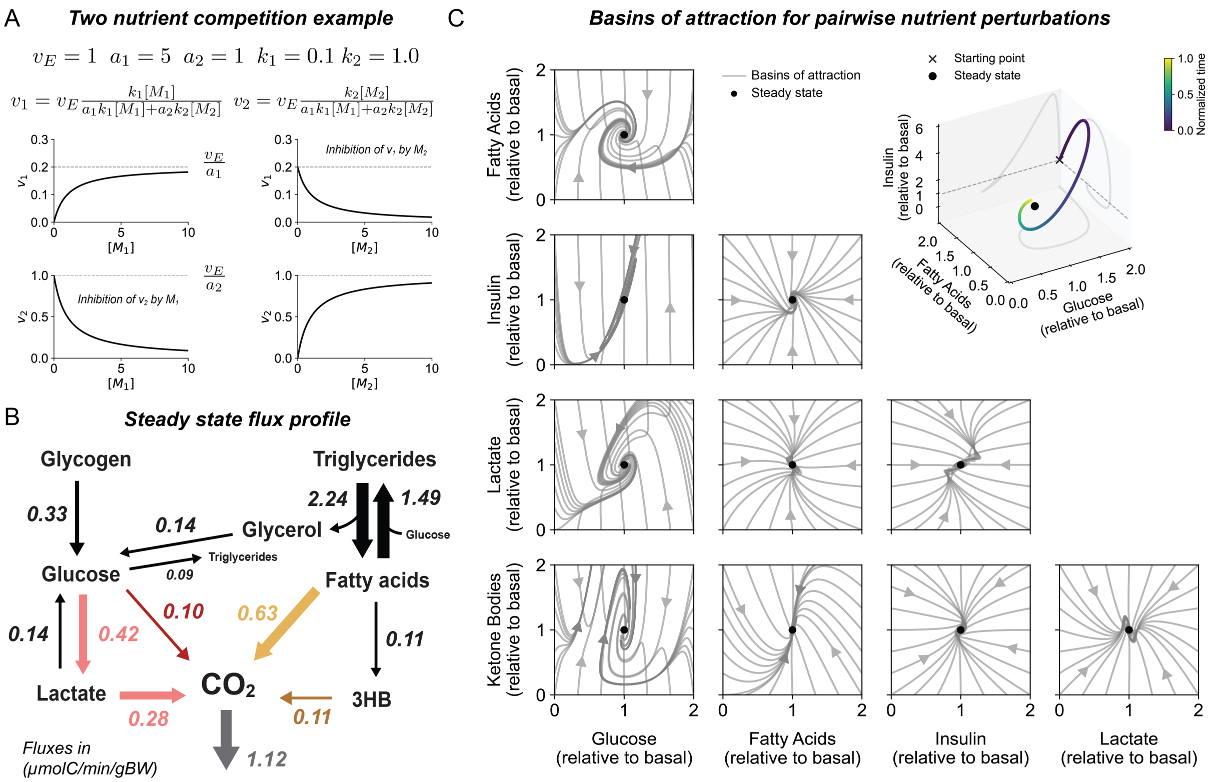


**Figure S1: The multi-nutrient model captures fasted physiology and is robust to nutrient and insulin perturbations, related to Figure 1.**

A) Example illustrating flux concentration relationships encoded by the competitive catabolism equation. Nutrients promote their own catabolism but inhibit the alternative substrates from being catabolized.

B) Steady-state fluxes used to derive mass action parameters of the multi nutrient model.

C) Basins of attraction for pairwise perturbations of fatty acids, glucose, lactate, 3HB and insulin.

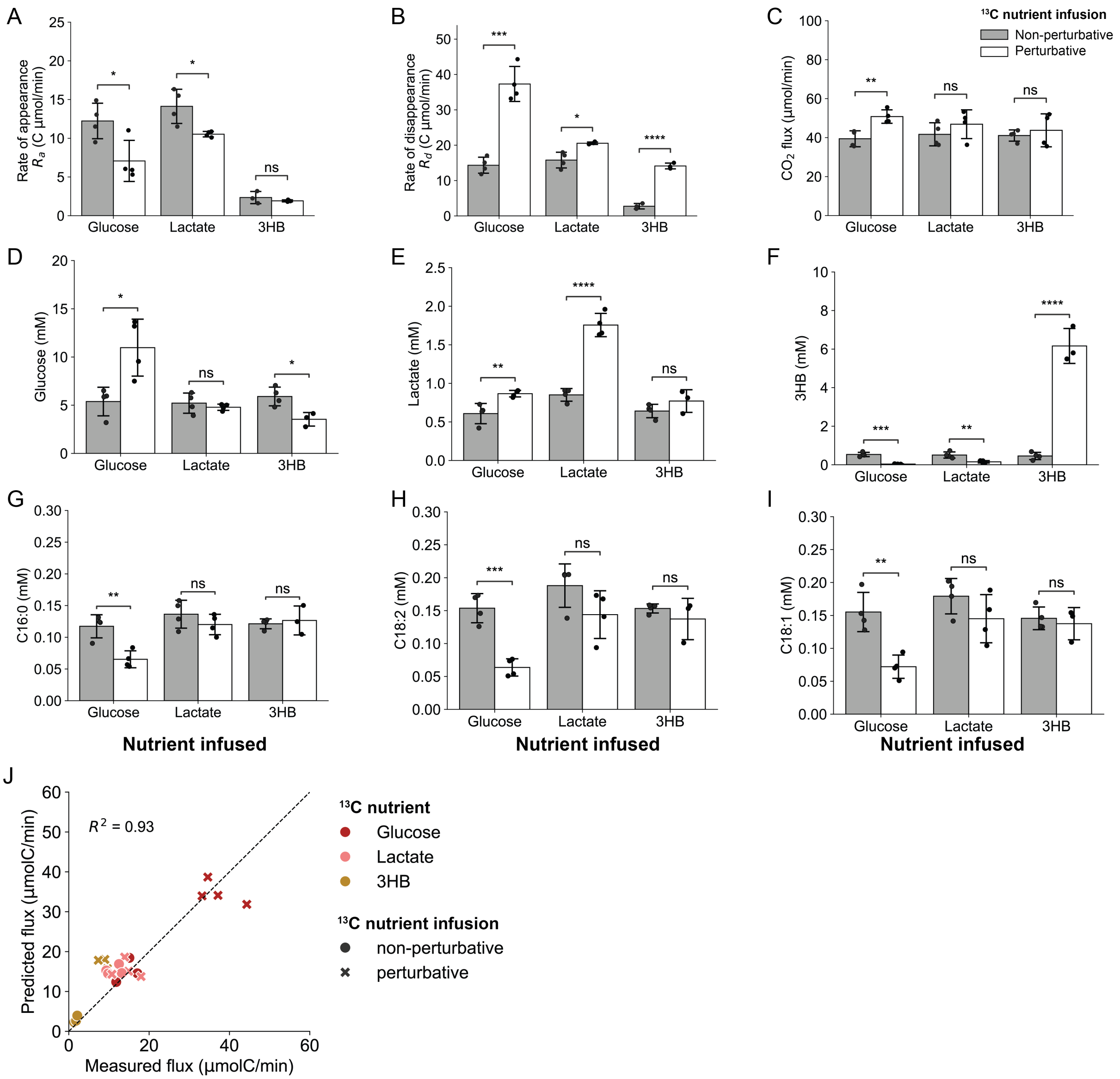


**Figure S2: Additional data from $^{13}$C-nutrient infusions, related to Figure 2.**

(A-C) Measured fluxes during intentionally perturbative or minimally perturbative nutrient infusions (n=3-4 mice per nutrient infusion): A) Endogenous production flux $R_a$, B) Total disposal flux $R_d$ and C) Flux of exhaled $CO_2$.

(D-I) Measured circulating concentrations during intentionally perturbative or minimally perturbative nutrient infusions (n=3-4 mice per nutrient infusion): D) Glucose, E) Lactate, F) 3HB, G) Palmitate (C16:0), H) Linoleate (C18:2), and I) Oleate (18:1).

J) Quantitative comparison of the competitive catabolism predictions and measured catabolic rates (related to Figure 2E,F).

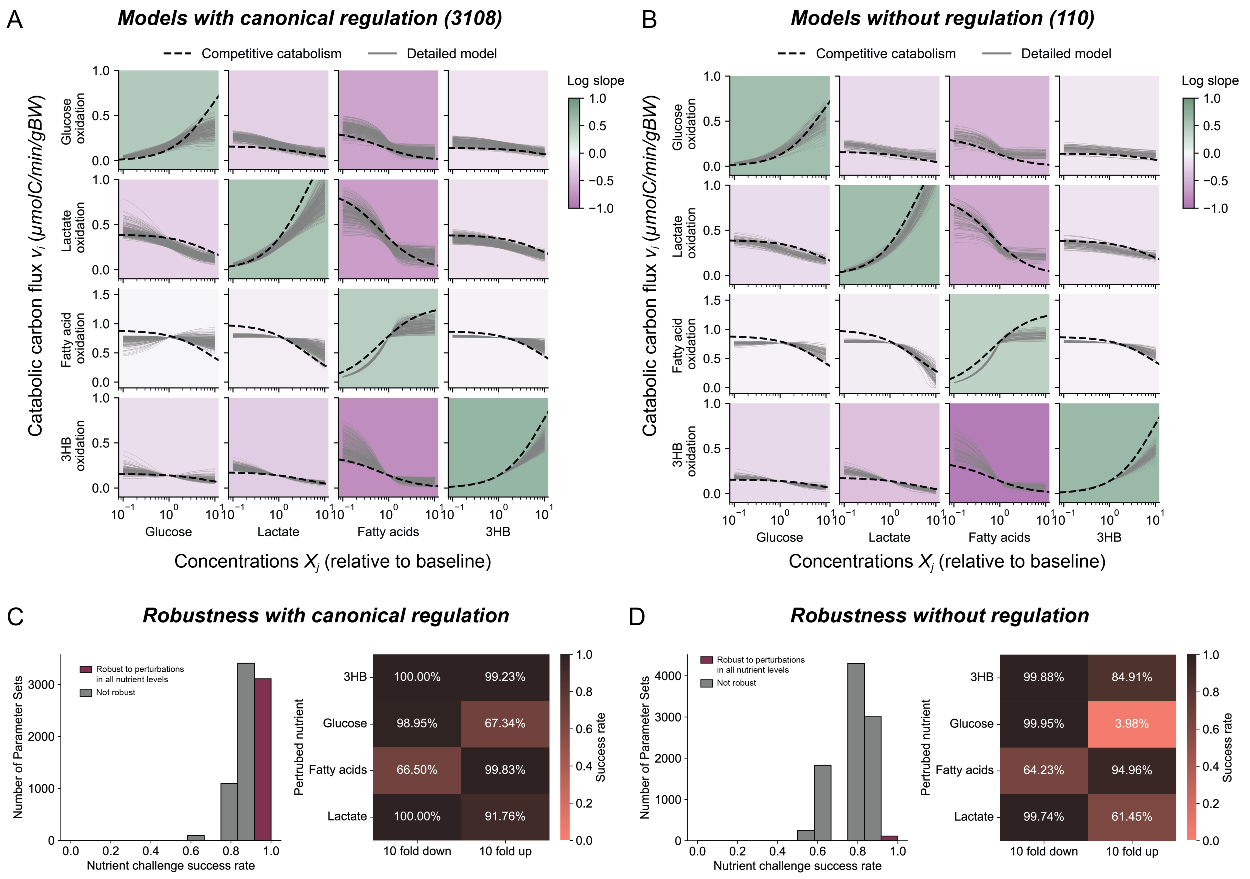


**Figure S3: Competitive catabolism also emerges in the absence of canonical regulation, related to Figure 3.**

(A-B) Simulated impact on metabolic flux of changing the concentration of the metabolite shown on the X-axis (green boxes, mass action; purple boxes, competitive catabolism). Solid gray lines represent outputs from the detailed kinetic model; dashed black lines indicate the competitive catabolism rate law: A) Models incorporating canonical competitive and allosteric regulation, (B) Models based solely Michaelis Menten like kinetics without regulation.

(C-D) Analysis of metabolic homeostatic capacity of the detailed kinetic model in response to changing nutrient availability evaluated across 20,000 sampled parameter sets: C) models including textbook cellular metabolic regulation, D) models based solely Michaelis Menten like kinetics without regulation. Canonical regulation results in greater robustness to nutrient perturbations.

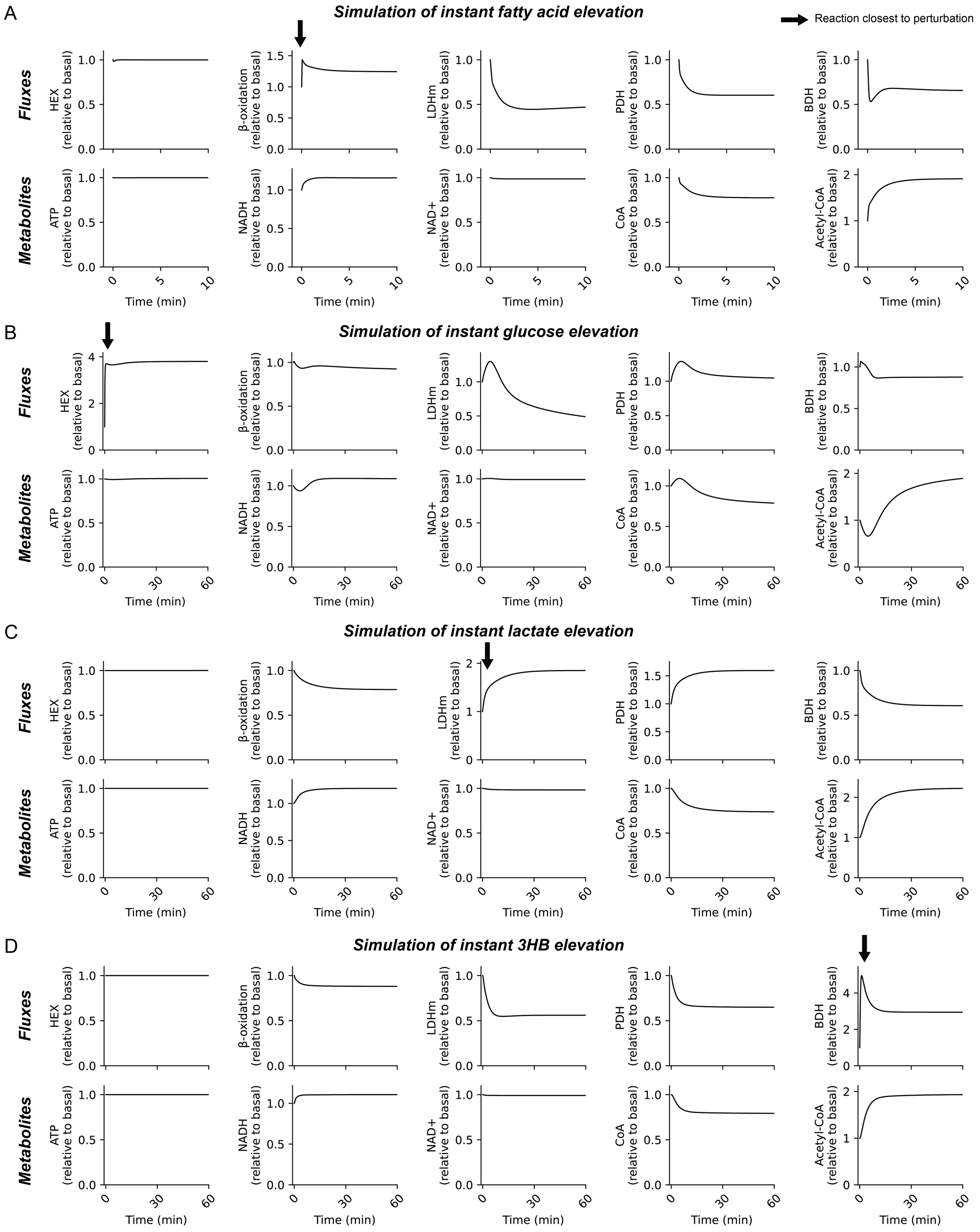


**Figure S4: NADH/NAD+ and Acetyl-CoA/CoA cofactor pairs are poised to mediate competition among nutrients.**

A) Fluxes and metabolite concentrations for dynamical simulations based on the detailed kinetic model for an instantaneous 5-fold increase in fatty acids.

B) Instantaneous 5-fold increase in glucose.

C) Instantaneous 5-fold increase in lactate.

D) Instantaneous 5-fold increase in 3HB.

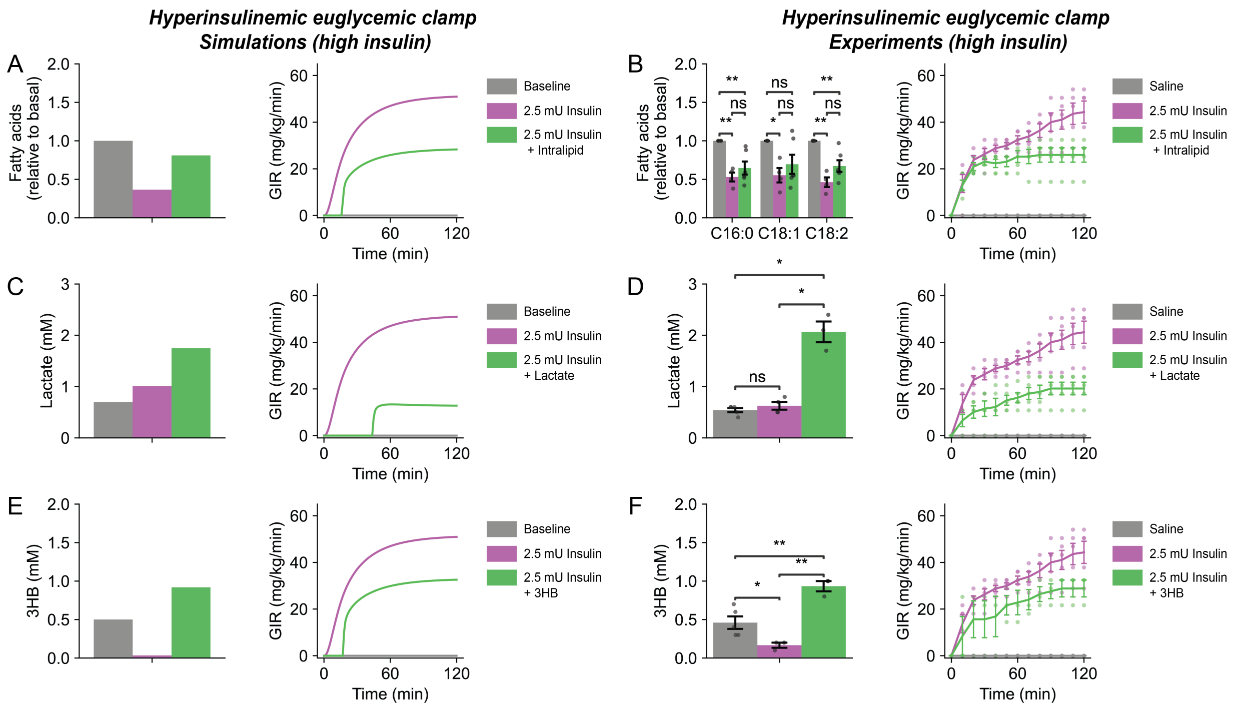


**Figure S5: Simulated and experimental euglycemic-hyperinsulinemic clamps at high insulin (2.5 mU), related to Figure 4.**

In addition to glucose infusion to maintain euglycemia, perturbative intralipid, lactate, or 3HB infusion were also infused as indicated. The exogenous nutrient infusions trigger competitive catabolism and thereby suppress insulin-induced glucose consumption. Simulations are directly from the multi-nutrient model in Fig. 1 and involve no fitted parameters. The same data for saline (n = 5 mice) and insulin (n = 4 mice) are repeated in multiple panels.

A) Simulation with intralipid infusion (simulated as fatty acid infusion).

B) Experiment with intralipid infusion (n=4 mice).

C) Simulation with lactate infusion.

D) Experiment with lactate infusion (n=3 mice).

E) Simulation with 3HB infusion.

F) Experiment with 3HB infusion (n=3 mice).

3HB, 3-hydroxybutyrate. ns, not significant. $*p < 0.05$, $**p < 0.01$, $***p < 0.001$, by two-sampled t-test.

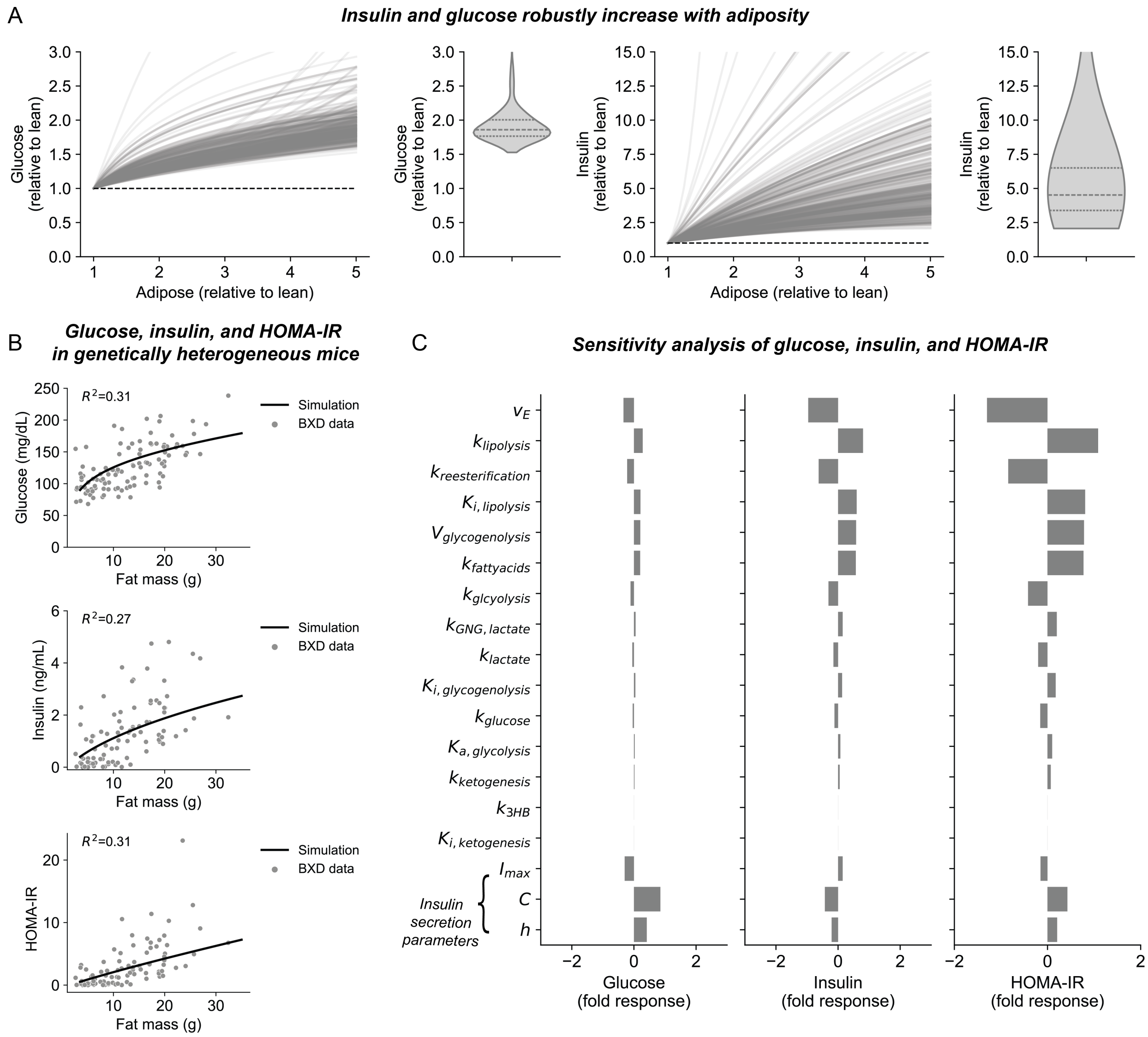


**Figure S6: Adiposity robustly induces hyperglycemia and hyperinsulinemia, related to Figure 6.**

A) Steady-state simulated glucose and insulin as a function of body fat for 500 models with randomly selected parameter values varying between 0.5 and 2 times the actual parameter value in the predictive multi-nutrient model.

B) Model predictions of glucose, insulin, and HOMA-IR as a function of body fat match experimental data from genetically heterogenous mice.

C) Parameter sensitivity analysis for glucose, insulin, and HOMA-IR in obesity. Positive values indicate increase with model parameters; negative values indicate inverse relationships. Parameters are listed in order of descending absolute sensitivity to insulin resistance (HOMA-IR). Parameters governing insulin secretion are listed separately at the bottom.

A

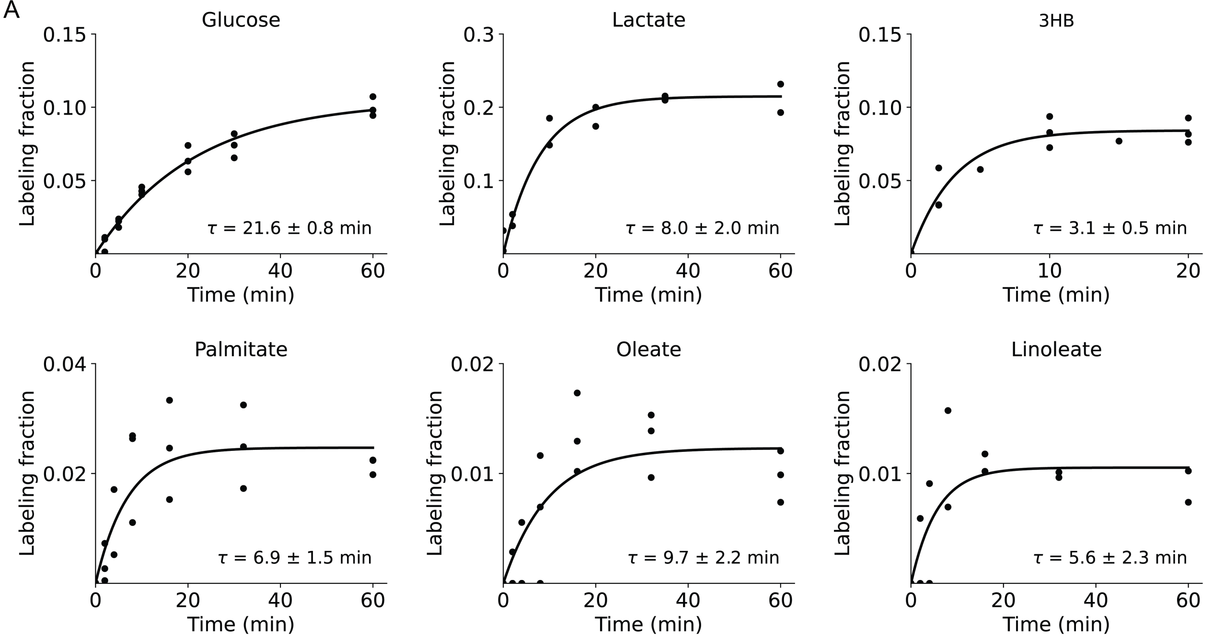


B

**Pool size**

$$P = F_{circ} \times \tau$$

**Volume of distribution**

$$V_d = \frac{P}{C}$$

C

| | $\tau$ (min) | $F_{circ}$* (nmol min$^{-1}$gBW$^{-1}$) | $P$ (µmol gBW$^{-1}$) | $C$ (mmol L$^{-1}$) | $V_d$ (mL gBW$^{-1}$) |
|---|---|---|---|---|---|
| *Glucose* | 21.6 | 95.4 | 2.0 | 6.0 | 0.33 |
| *Lactate* | 8.0 | 151.7 | 1.2 | 0.7 | 1.7 |
| *3HB* | 3.1 | 27.5 | 0.09 | 0.5 | 0.18 |
| *Palmitate* | 6.9 | 24.0 | 0.17 | 0.12 | 1.42 |
| *Oleate* | 9.7 | 29.5 | 0.29 | 0.16 | 1.81 |
| *Linoleate* | 5.6 | 47.6 | 0.27 | 0.16 | 1.68 |
| *Fatty acids*** *(overall pool)* | 7.5 | 126.4 | 0.94 | 0.5 | 1.88 |

**Figure S7: Determining volumes of distribution for the metabolites in the multi-nutrient model**
(A) Kinetics of tracer enrichment in blood during infusion of $^{13}$C-labeld nutrients (Glucose, Lactate, 3HB, Palmitate, Oleat, and Linoleate). (B) Relevant equations to compute volumes of distribution from labeling time constants, circulatory flux, and blood concentrations. (C) Table with pool size and volume of distribution calculations for modeled metabolites.

*Circulatory fluxes were measure previously[92]

**Overall fatty acid pool, $\tau$ was computed from the abundance weighted average of the three fatty acids, and $F_{circ}$ estimated assuming together Palmitate, Oleate and Linoleate account for 80% of the total circulating fatty acids (20% Palmitate, 30% Oleate, and 30% Linoleate).

3HB, 3-hydroxybutyrate.

**Table S1: Mathematical expressions for fluxes in the multi-nutrient model**

| Rate name | Symbol | Expression | Description |
|---|---|---|---|
| Insulin secretion | $v_{I,0}$ | $I_{max}\left(\frac{[G]^h}{C^h+[G]^h}\right)$ | Hill kinetics secretion of insulin in response to glucose |
| Insulin degradation | $v_{I,d}$ | $k_d[I]$ | Degradation of circulating insulin by mass action |
| Insulin receptor detection | $v_{IR,0}$ | $\frac{[I]}{\tau}$ | Activation rate of insulin action by circulating insulin |
| Insulin receptor off rate | $v_{IR,d}$ | $\frac{[I_A]}{\tau}$ | Deactivation rate of insulin action by mass action |
| Glycogenolysis | $v_{glycogenolysis}$ | $V_{glycogenolysis}\left(1-\frac{[I_A]}{[I_A]+K_{i,glycogenolysis}}\right)$ | Zeroth order glycogen degradation inhibited by insulin action, assumes constant glycogen pool. |
| Lipolysis | $v_{lipolysis}$ | $k_{lipolysis}[FM]\left(1-\frac{[I_A]}{[I_A]+K_{i,lipolysis}}\right)$ | Fat mass action inhibited by insulin action. |
| Reesterification | $v_{reesterification}$ | $k_{reesterification}[F]$ | Fatty acid mass action |
| Gluconeogenesis | $v_{GNG,lactate}$ | $k_{GNG,lactate}[L]$ | Lactate mass action |
| Ketogenesis | $v_{ketogenesis}$ | $k_{ketogenesis}[F]\left(1-\frac{[I_A]}{[I_A]+K_{i,ketogenesis}}\right)$ | Fatty acid mass action inhibited by insulin action. |
| Glycolysis | $v_{glycolysis}$ | $\frac{k_{glycolysis}[G]\left(1+a\frac{[I_A]}{[I_A]+K_{a,glycolysis}}\right)}{D}$ | Competitive catabolism for glycolysis, contains additional activation of glycolysis by insulin. |
| Glucose oxidation | $v_{glucose}$ | $\frac{k_{gluose}[G]}{D}$ | Competitive catabolism for direct glucose oxidation |
| Lactate oxidation | $v_{lactate}$ | $\frac{k_{lactate}[L]}{D}$ | Competitive catabolism for lactate oxidation |
| Fatty acid oxidation | $v_{fatty\ acids}$ | $\frac{k_{fatty\ acids}[F]}{D}$ | Competitive catabolism for fatty acid oxidation |
| 3HB oxidation | $v_{3HB}$ | $\frac{k_{3HB}[K]}{D}$ | Competitive catabolism for 3HB |
| Common Denominator | $D$ | $k_{glycolysis}[G]\left(1+a\frac{[I_A]}{[I_A]+K_{a,glycolysis}}\right)$ $+k_{lactate}[L]+\ k_{fatty\ acids}[F]+k_{glucose}[G]$ $+\ k_{3HB}\ [K]$ | Common denominator for all competitive catabolism expressions |

### Table S2: Steady-state concentrations, fluxes and parameters for the multi-nutrient model

| | Symbol | Dimension less val. | Unit | Value |
|---|---|---|---|---|
| **Steady state levels** | | | | |
| Circulating insulin | $[I_0]$ | 0.055 | ng mL$^{-1}$ | 0.4 |
| Insulin action | $[I_{A,0}]$ | 0.055 | ng mL$^{-1}$ | 0.4 |
| Glucose | $[G_0]$ | 1.0 | mM | 7 |
| Lactate | $[L_0]$ | 1.0 | mM | 0.7 |
| 3HB | $[K_0]$ | 1.0 | mM | 0.5 |
| Fatty acids | $[F_0]$ | 1.0 | mM | 0.5 |
| **Steady state fluxes** | | | | |
| Glycogenolysis | $v_{glycogenolysis}$ | 0.0080 | nmol min$^{-1}$ gBW$^{-1}$ | 56 |
| Lipolysis | $v_{lipolysis}$ | 0.0067 | nmol min$^{-1}$ gBW$^{-1}$ | 47 |
| Reesterification | $v_{reesterification}$ | 0.0044 | nmol min$^{-1}$ gBW$^{-1}$ | 31 |
| Gluconeogenesis | $v_{GNG,lactate}$ | 0.0033 | nmol min$^{-1}$ gBW$^{-1}$ | 23 |
| Ketogenesis | $v_{ketogenesis}$ | 0.0010 | nmol min$^{-1}$ gBW$^{-1}$ | 7 |
| Glycolysis | $v_{glycolysis}$ | 0.0100 | nmol min$^{-1}$ gBW$^{-1}$ | 70 |
| Glucose oxidation | $v_{glucose}$ | 0.0024 | nmol min$^{-1}$ gBW$^{-1}$ | 17 |
| Lactate oxidation | $v_{lactate}$ | 0.0133 | nmol min$^{-1}$ gBW$^{-1}$ | 93 |
| Fatty acid oxidation | $v_{fatty\ acids}$ | 0.0057 | nmol min$^{-1}$ gBW$^{-1}$ | 39 |
| Energy demand (ATP) | $v_E$ | 1.0 | μmol min$^{-1}$ gBW$^{-1}$ | 7 |
| **Parameters** | | | | |
| Fat mass relative to lean | $[FM]$ | 1.0 | - | 1.0 |
| Maximal Insulin secretion rate | $I_{max}$ | 1.0 | ng mL$^{-1}$ min$^{-1}$ | 0.2 |
| Half maximal glucose concentration | $C$ | 2.3 | mM | 16 |
| Hill coefficient | $h$ | 3.4 | - | 3.4 |
| Insulin degradation rate | $k_d$ | 1.0 | min$^{-1}$ | 0.5 |
| Insulin action delay | $\tau$ | 1.0 | min | 30 |
| Volume of distribution of glucose | $V_G$ | - | mL gBW$^{-1}$ | 0.33 |

| | | | | |
|---|---|---|---|---|
| Volume of distribution of lactate | $V_L$ | - | mL $gBW^{-1}$ | 1.7 |
| Volume of distribution of fatty acids | $V_F$ | - | mL $gBW^{-1}$ | 1.88 |
| Volume of distribution of 3HB | $V_K$ | - | mL $gBW^{-1}$ | 0.18 |
| Affinity constant for insulin inhibition of ketogenesis | $K_{i,ketogenesis}$ | 0.1 | | |
| Affinity constant for insulin inhibition of lipolysis | $K_{i,lipolysis}$ | 1 | | |
| Affinity constant for insulin inhibition of glycogenolysis | $K_{i,glycogenolysis}$ | 10 | | |
| Affinity constant for insulin activation of glycolysis | $K_{a,glycolysis}$ | 10 | mg $dL^{-1}$ | 230 |
| Extent of activation for glycolysis[93] | $a$ | 2.5 | - | 2.5 |

**Table S3 Constrained model parameters for the multi-nutrient model**

| Parameter name | Symbol | Expression |
|---|---|---|
| Glycogen breakdown $V_{max}$ | $V_{glycogenolysis}$ | $\frac{v_{glycogenolysis}}{1-\frac{[I_{A,0}]}{[I_{A,0}]+K_{i,glycogenolysis}}}$ |
| Rate constant lactate gluconeogenesis | $k_{GNG,lactate}$ | $\frac{v_{GNG,lactate}}{[L_0]}$ |
| Rate constant reesterification | $k_{reesterification}$ | $\frac{v_{reesterification}}{[F_0]}$ |
| Rate constant lipolysis | $k_{lipolyis}$ | $\frac{v_{lipolysis}}{A\left(1-\frac{[I_{A,0}]}{[I_{A,0}]+K_{i,lipolysis}}\right)}$ |
| Rate constant ketogenesis | $k_{ketogenesis}$ | $\frac{v_{ketogenesis}}{[F_0]\left(1-\frac{[I_{A,0}]}{[I_{A,0}]+K_{i,glycogenolysis}}\right)}$ |
| Rate constant glycolysis | $k_{glycolysis}$ | $\frac{v_{glycolysis}}{[G_0](1+a\frac{[I_{A,0}]}{[I_{A,0}]+K_{a,glycolysis}})}$ |
| Rate constant glucose oxidation | $k_{glucose}$ | $\frac{v_{glucose}}{[G_0]}$ |
| Rate constant lactate oxidation | $k_{lactate}$ | $\frac{v_{lactate}}{[L_0]}$ |
| Rate constant fatty acids oxidation | $k_{fatty\ acids}$ | $\frac{v_{fatty\ acids}}{[F_0]}$ |
| Rate constant 3HB oxidation | $k_{3HB}$ | $\frac{v_{3HB}}{[K_0]}$ |
